# Diversification of functional requirements for proteolysis of Auxin Response Factors

**DOI:** 10.1101/2025.09.03.673984

**Authors:** Martijn de Roij, Esmée Heijdra, Ryuichi Nishihama, Jan Willem Borst, Dolf Weijers

## Abstract

Auxin signaling through the Nuclear Auxin Pathway is essential for plant development and is mediated by competing A-class and B-class Auxin Response Factor (ARF) transcription factors. Recently, proteasomal ARF degradation through a degradation signal embedded within the DNA-Binding Domain, was identified as a key component of auxin response. Here, we investigate the structural requirements and biological relevance of ARF degradation in the bryophyte *Marchantia polymorpha*. We identify a critical residue for proteolysis of the repressive, B-class MpARF2, and find it to be functionally conserved in degradation of the activating, A-class MpARF1. Unlike MpARF2, however, impaired MpARF1 degradation had little effect on auxin response and development, suggesting differential integration in biological function. We find that MpARF2 degradation occurs across all developmental stages of the life cycle, and is required for MpARF2 function during development. Our findings reveal a degradation mechanism targeting A- and B-class ARFs which shares the same origin, but has evolved along unique evolutionary trajectories.

## Introduction

Auxin is an important signaling molecule in plant development as it orchestrates several physiological processes^1^. While multiple mechanisms through which auxin elicits these responses have been revealed, the best characterized auxin mode of action to date relies on the control of gene transcription through the Nuclear Auxin Pathway (NAP)^2,3^. The NAP revolves around three families proteins, TRANSPORT INHIBITOR RESPONSE1/AUXIN SIGNALING F-BOX proteins (TIR1/AFBs), auxin/indole-3-acetic acid (Aux/IAA), and AUXIN RESPONSE FACTORS (ARFs)^4^. TIR1/AFBs are components of a SKP-CULLIN-F-BOX ubiquitin-ligase complex (SCF^TIR/AFB^) and form a coreceptor complex together with Aux/IAA proteins in presence of auxin, which leads to ubiquitylation and subsequent degradation of Aux/IAAs by the proteasome^5-7^. Given the role of Aux/IAAs as ARFs inhibitors, their degradation subsequently releases the ARFs from an inhibitory state leading to transcriptional activation of auxin-responsive genes^8,9^. ARFs are central in this system as they connect perception of auxin by TIR1/AFBs and Aux/IAAs to a transcriptional response which ultimately leads to physiological changes.

In land plants, ARFs contain an N-terminal DNA-binding domain (DBD), a middle region (MR), and a Phox and Bem 1 domain (PB1), and can be classified into three main subfamilies^10^. Two of these families, the A- and B-class ARFs, possess a highly conserved DNA-binding domain which show a substantial overlap in functionality and binding specificity of their target genes^11-13^. However, A-class ARFs tend to function as auxin-induced gene activators, whereas B-class ARFs are auxin-independent repressors^13,14^. A- and B-Class ARFs compete for DNA-binding sites in a stoichiometry-dependent way^13,14^, and the relative abundance of A- and B-class ARFs within the nucleus therefore dictates cellular transcriptional responsiveness to auxin.

To maintain an appropriate ARF stoichiometry, plants rely on both transcriptional and post translational control mechanisms^15,16^. In recent years, ARF proteolysis through the 26S proteasome has emerged as a critical node in auxin signaling^17-19^. Though various ARFs are known to be unstable, the precise molecular mechanisms and biological importance remain poorly understood^15^.

Recent findings identified a conserved motif in the DBD that is critical for their proteasome-mediated degradation across diverse plant lineages^20,21^. Mutation of residues in this motif led to B-class ARF stabilization, which in turn represses the auxin response, ultimately leading to impaired development^20,21^. In the bryophyte *Marchantia polymorpha* (hereafter Marchantia), each major ARF family is represented by a single gene copy, making it an ideal, minimal system to dissect core principles of auxin signaling^22^. The sole B-class ARF MpARF2 carries the conserved degradation motif, and mutations in residues comprising this motif severely impacted normal development^21^. Swapping this motif of MpARF2 with homologous motifs of other ARFs, representing all known ARF classes across the green lineage, revealed that the instability conferred by this motif likely emerged in an ancestral AB-ARF that gave rise to the A- and B-class ARF clades^21-23^. While the importance of degradation through this motif was demonstrated in B-class ARFs, it remains unknown if this instability was integrated in A-class ARF function. Furthermore, many aspects of B-class ARF degradation, such as structural requirements, spatiotemporal patterns and biological relevance, remain unresolved.

Here, we leveraged Marchantia as a model to obtain deeper insight into the molecular determinants regulating ARF proteolysis. Our work demonstrates how precise dosing of ARF levels through proteasome-mediated degradation contributes to auxin response and development in land plants.

## Results

### Structural requirements of MpARF2 degradation

We have previously demonstrated that a small loop in the MpARF2 DBD is required for regulated protein degradation. Single and double mutations in the E297 (E297K – a charge reversal) and R300 (R300Q – a loss of charge) residues radically decrease protein turnover, with severe consequences for development^21^. The mutations that were previously engineered were inspired by induced mutations in *Physcomitrium patens* ARFs that caused auxin-related phenotypes^20^. However, given that no systematic analysis of the degradation region in any ARF has yet been reported, it is unclear what the requirements of this region for effective degradation are. Particularly because both mutations lead to a loss or reversal of charge, it is possible that this charge is critical for protein degradation. There is substantial variation in the degradation motif among ARFs, and the unstable maize ZmARF28 lacks the charged residues, urging the question of what properties underlie protein instability.

To address such questions, we performed a systematic mutagenesis scan by replacing each residue flanking the E297 and R300 amino acids in MpARF2 with an alanine, except A294 and A295, which were substituted by a glycine (Fig. 1, A). None of these mutations had a strong predicted effect on overall MpARF2 DBD folding (fig. S1, A). We generated transgenic plants expressing the MpARF2 DBD fused to mNeonGreen (mNG) under control of the native Mp*ARF2* promoter in the Tak-1 background. We used only the DBD to prevent phenotypic effects, and because we previously found that remaining MpARF2 domains did not contribute to protein stability^21^. We next quantified fluorescence intensity in nuclei of transgenic gemmae as a proxy for protein stability. Although not statistically significant, the E297A, K298A, and F301A mutations all seemed to have a very weak stabilizing effect, whereas A294G, A295G, T296A, and S299A behaved identically to the wild type (WT) protein, indicating that none of these residues are critical for protein degradation (Fig. 1, A and B). In addition, we treated all lines with the proteasome inhibitor Bortezomib (Bz), and found that each accumulated, excluding the possibility that low protein accumulation observed was due to low expression levels (fig. S1, B).

**Fig. 1.**
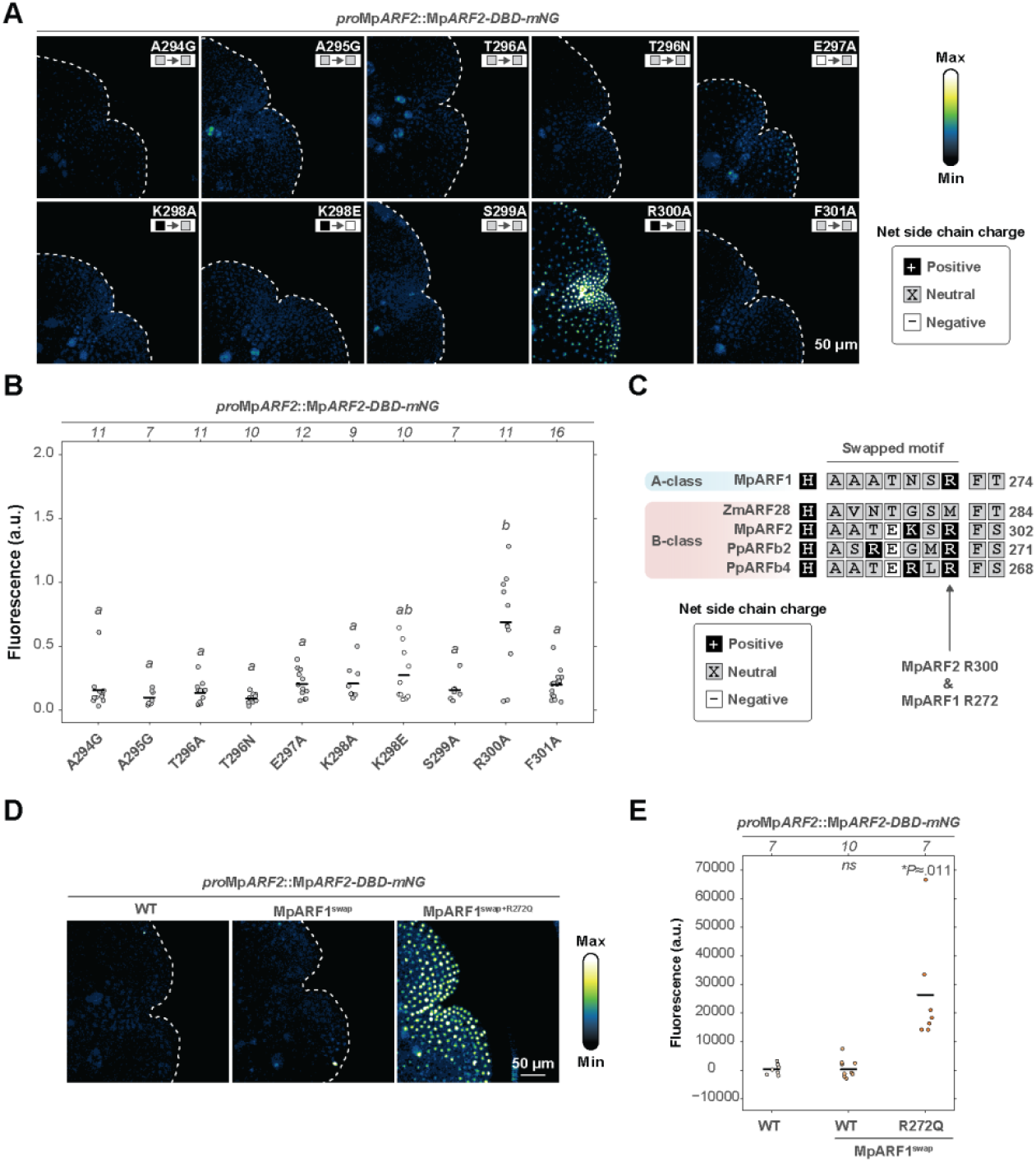
A positive charge is essential for MpARF degradation. (**A**) Representative confocal images showing fluorescence patterns in dormant gemmae expressing an *proMpARF2*::Mp*ARF2*-*DBD*-*mNeonGreen* fusion with indicated mutations. Amino acid side chain charge is indicated. (**B**) Quantification of overall fluorescence in several independent transgenic lines of which one is shown in (A). Letters in italics indicate significant differences determined per one-way ANOVA (*F*=(9,94)=6.322, *P≈*5.85×10^-7^) and Tukey HSD post hoc test with Bonferroni correction (*P*<.05). (**C**) Protein alignment of a selection of ARF proteins, amino acid side chain charge and the swapped motif are indicated. (**D**) Representative confocal images of *pro*Mp*ARF2*::Mp*ARF2*-*DBD*-*mNeonGreen* fusions with indicated motif swaps. (**E**) Quantification of (D), statistical differences from the WT were determined per T-test (*P*<.05). (B and C) The sample size is indicated above the figure, each replicate consists of a gemmae from an independent transgenic line. (B and E) Exact *P*-values are provided in the Source dataset.

Intriguingly, in contrast to the E297K mutation^21^, E297A fluorescence levels remained low (Fig. 1, A and B), suggesting that the charge reversal, rather than loss of negative charge conferred E297K stability. The R300A mutation did induce protein stability (Fig. 1, A and B), hence loss of positive charge, rather than specific substitution to Q, caused R300Q stabilization. Collectively, these observations suggest that removal of the R300 positive charge or gain of E297K positive charge has a dramatic impact on protein instability. We therefore engineered a charge reversal in K298 (K298E), and found that this weakly stabilized the protein (Fig. 1, A and B), suggesting that interactions mediating protein degradation are likely driven by charges, yet these act in a localized manner.

In the Maize ZmARF28 protein, a mutation equivalent to MpARF2 S299N causes stabilization^20^, but we previously found this mutation in MpARF2 not to have similar effects (Fig. 1, C)^21^. Given the existance of one additional alcoholic residue in the MpARF2 degradation motif (T296), we mutated this threonine to asparagine (T296N) to test if the N side chain may influence stability in a different position in MpARF2. The T296N mutation did not affect protein stability (Fig. 1, A and B). This suggests that the degradation motif in MpARF2 is lenient to mutation, but depends on positive charges, most prominently R300.

The key single R300 degradation residue in MpARF2 is conserved in the A-class ARF MpARF1^21^, indicative of a possibly conserved proteolytic degradation mechanism shared by the two classes (Fig. 1, C). We previously swapped the equivalent region from MpARF1 into the MpARF2 DBD, and observed conferred instability^21^. To test if this instability required the conserved arginine, we engineered the R272Q mutation within the MpARF2 DBD^MpARF1 swap^ fusion protein, and found this to have a drastic, stabilizing effect (Fig 1, C and D; fig. S1, C). Thus, protein degradation in Marchantia MpARF1 and MpARF2 may require a minimal arginine, and may be mediated by the same mechanism.

### Regulated degradation of MpARF1

We previously hypothesized that proteasome-mediated proteolysis through the ARF DBD emerged in an ancestral AB-class ARF protein^21^. However, when following the accumulation of MpARF1 and MpARF2 in the same plant over time, either in presence or absence of a proteasome inhibitor, we found that dynamics of accumulation are rather different: proteasome inhibition leads to excess MpARF2 accumulation, whereas MpARF1 stops being degraded, but does not accumulate beyond regular levels^18^. It is therefore a question whether MpARF1 is actively degraded.

Given that the MpARF1 R272Q mutation stabilized MpARF2 when introduced as part of a region swap (Fig. 1, D), we first engineered the same mutation in the full length MpARF1 protein (MpARF1-FL), and expressed this as an mNG fusion driven by the native Mp*ARF1* promoter (Fig. 2, A). We found this mutation to weakly, but statistically significant, stabilize the MpARF1 protein (Fig. 2, B and C). Notably, Bz treatment led to increased accumulation of MpARF1-FL (Fig. 2, D to F). Next, to determine which protein domains confer MpARF1 instability, we expressed full length wild type (MpARF1-FL^WT^) and R272Q MpARF1 (MpARF-FL^R272Q^), as well as individual domains (MpARF1-DBD^WT^ and MpARF1-DBD^R272Q^, MpARF1-MR, and MpARF1-PB1) as mNG fusions, which were expressed from the constitutive *UBIQUITIN-CONJUGATING ENZYME E2* promoter (*pro*Mp*UBE2)* to prevent autoregulation of the Mp*ARF1* promoter^24^. Interestingly, we found that MpARF1-FL accumulated at lower levels than each of its separate domains, indicative of a collective contribution to protein instability (fig. S2, A and B). When expressed from the Mp*UBE2* promoter, we did not find a significant effect of the R272Q mutation on protein (domain) stability (fig. S2, A and B). Fluorescence levels varied greatly between transgenics, particularly for the DBD-only lines, which may have occluded effects of the mutation (fig. S2, A and B). Likewise, we observed the same variation when expressing the MpARF1-DBD from the Mp*ARF2* promoter (Fig S2, C and D). We therefore tested the effect of Bz treatment on *pro*Mp*UBE2*::Mp*ARF1*-*DBD*^*WT*^ lines and found this to cause increased accumulation (Fig 2, G to I). We next observed protein accumulation over time following germination of gemmae in *pro*Mp*ARF1*::Mp*ARF*-*FL*-*mNG* and *pro*Mp*UBE2*::Mp*ARF1*-*DBD*-*mNG* lines in which the R272Q was engineered. In both these contexts, we found decline of the WT protein over time, and reduced decline induced by the R272Q mutation, across the meristematic region (Fig. 3, A to F; fig. S2, E). Thus, we conclude that the MpARF1 DBD confers instability, likely through the conserved R272.

**Fig. 2.**
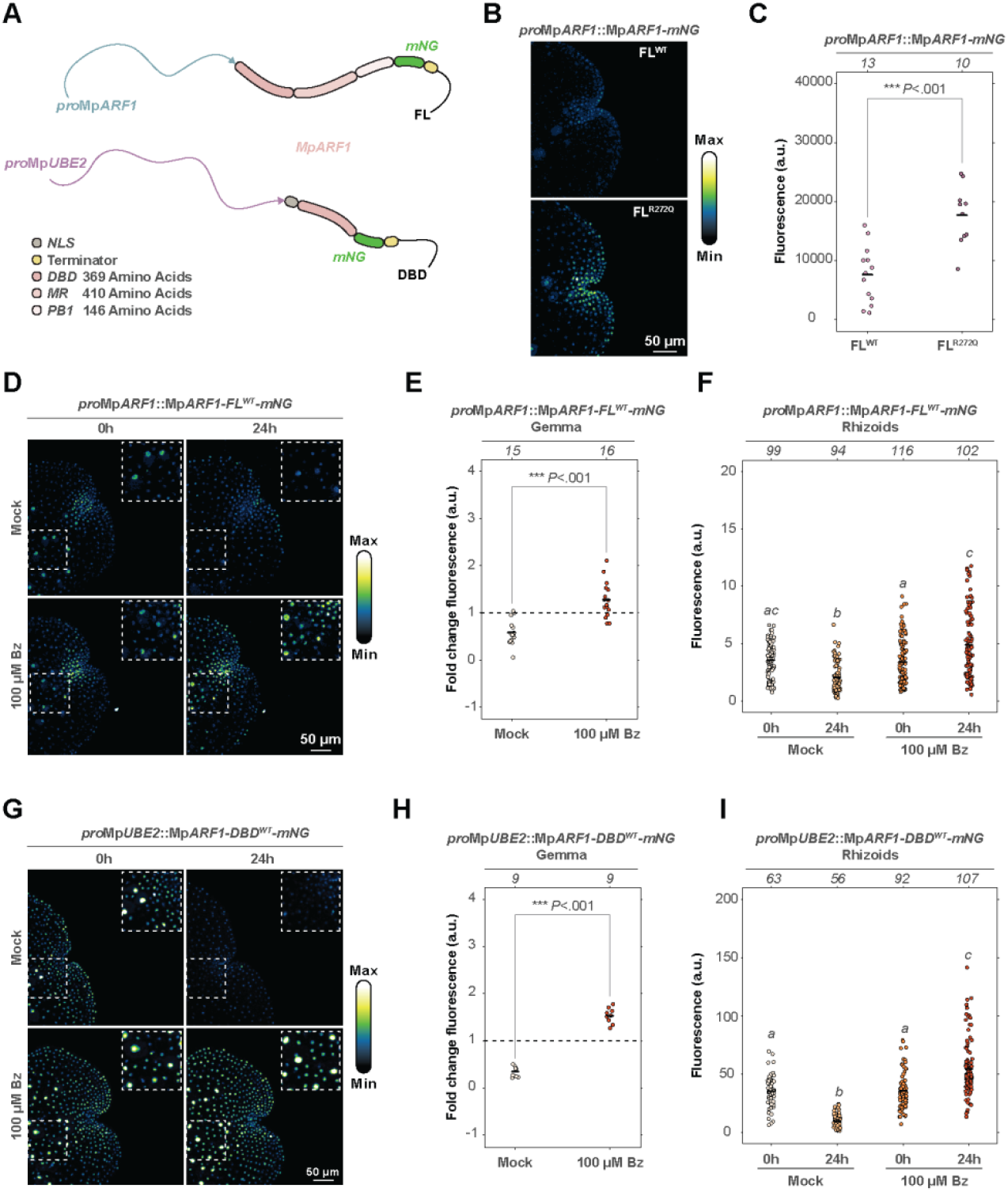
Requirements for MpARF1 proteolysis. (**A**) Diagram depicting genetic constructs encoding translational fusions to mNeonGreen (mNG) analyzed in this experiment; names correspond to other panels in this figure. (**B**) Representative confocal images showcasing *proMpARF1*::Mp*ARF1*-*FL*-*mNG* accumulation patterns in dormant gemmae. (**C**) Fluorescence intensity quantification of dormant gemmae expressing *pro*Mp*ARF1*::Mp*ARF1*-*FL*-*mNG*, both a wild type (WT) and R272Q mutant (R272Q) version, of which one is shown in (B). Fluorescence intensity was compared via T-test (*P*<.05). (**D**) Representative confocal images showing *pro*Mp*ARF1*::Mp*ARF1*-*FL*^*WT*^- *mNG* accumulation patterns in gemmae treated with Bortezomib (Bz) or Mock (DMSO) for 24 hours. The outlined portion of the image is enlarged. (**E**) Quantification of fold change in overall fluorescence of *pro*Mp*ARF1*::Mp*ARF1*-*FL*^*WT*^-*mNG* expressing gemmae, of which representatives are shown in (D), after 24 hours of treatment with Bortezomib (Bz) or Mock (DMSO). Fold change was compared per T-test (*P*<.05). (**F**) Mean fluorescence measured in individual nuclei of rhizoid initial cells before and after 24 hours of treatment with Bortezomib (Bz) or Mock (DMSO). Intensities were compared via Kruskal-Wallis test (*H*(3)=74.539, *P*≈4.548×10^-16^) with a Dunn post hoc test (*P*<.05). Same experiment as (D and E). (**G**) Representative confocal images showing *pro*Mp*UBE2*::Mp*ARF1*-*DBD*^*WT*^-*mNG* accumulation patterns in gemmae treated with Bortezomib (Bz) or Mock (DMSO) for 24 hours. An outlined portion of the image is enlarged. (**H**) Quantification of fold change in overall fluorescence of *pro*Mp*UBE2*::Mp*ARF1*-*DBD*^*WT*^-*mNG* expressing gemmae of which representatives are shown in (G) after 24 hours of treatment with Bortezomib (Bz) or Mock (DMSO). Fold change was compared per T-test (*P*<.05). (**I**) Mean fluorescence measured in individual nuclei of rhizoid initial cells before and after 24 hours of treatment with Bortezomib (Bz) or Mock (DMSO). Intensities were compared via Kruskal-Wallis test (*H*(3)=170.32, *P≈*2.2×10^-16^) with a Dunn post hoc test (*P*<.05). Same experiment as (G and H). For (C), (E), and (H): sample size is indicated above the figures, each replicate consists of the average of a gemmae representing a total of three independent transgenic lines. For (F), and (I): sample size is indicate above the figure, each replicate consists of a measurement from an individual nucleus of a rhizoid initial cell. (C, E-F, H-I) Exact *P*-values are provided in the Source dataset.

**Fig. 3.**
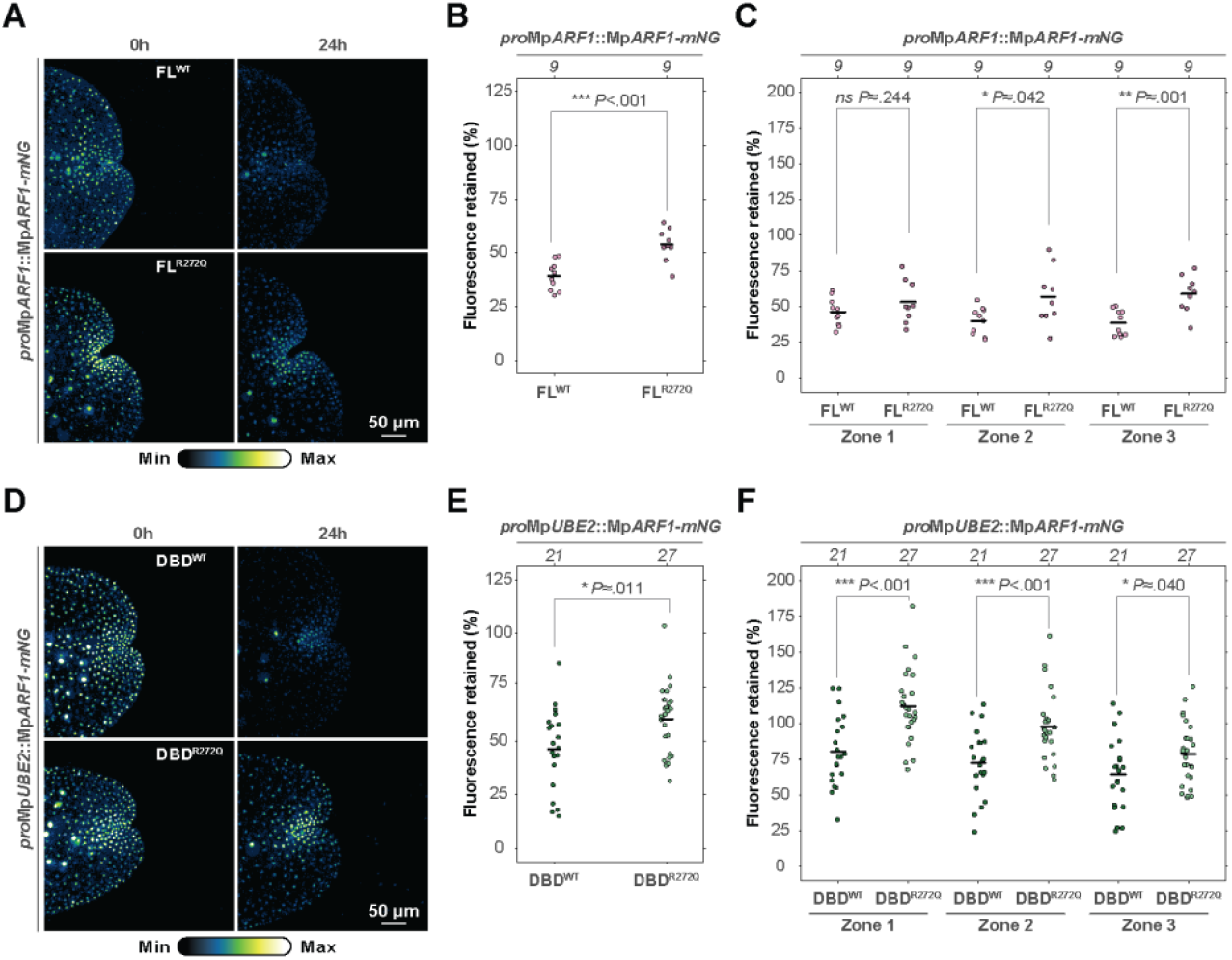
The R272Q mutation stabilizes MpARF1. (**A**) Representative confocal images showing *pro*Mp*ARF1*::Mp*ARF1*-*FL*-*mNG* accumulation patterns in individual gemmae in a dormant state (0 hours) and after germination (24 hours). (**B**) Quantification of the overall level of fluorescence retained in *pro*Mp*ARF1*::Mp*ARF1*-*FL*-*mNG* after 24 hours of germination, compared by T-test (*P*<.05). (**C**) Quantification of the level of fluorescence retained in three different zones of the apical notch, with Zone 1 corresponding to the zone most proximal to the apical cell, and Zone 3 being most distal of the apical cell, as illustrated in fig. S2, E. Fluorescence retained was compared by T-test (*P*<.05). Same experiment as (A and B). (**D**) Representative confocal images showing *pro*Mp*UBE2*::Mp*ARF1*-*DBD*^*WT*^-*mNG* accumulation patterns in individual gemmae in a dormant state (0 hours) and after germination (24 hours). (**E**) Quantification of the overall level of fluorescence retained in *pro*Mp*UBE2*::Mp*ARF1*-*DBD*^*WT*^-*mNG* after 24 hours of germination, compared by T-test (*P*<.05). (**F**) Quantification of the level of fluorescence retained in three different zones of the apical notch, with Zone 1 corresponding to the zone most proximal to the apical cell, and Zone 3 being most distal of the apical cell, as illustrated in Fig. S2, E. Fluorescence retained was compared by T-test (*P*<.05). Same experiment as (D and E). (B and C) Sample size is indicated above the figures, each replicate consists of the average of a gemmae (n=3) representing three independent transgenic lines in total. (E and F) Sample size is indicated above the figures, each replicate consists of the average of a gemmae (n=7-9) representing three independent transgenic lines in total. (B-C and E-F) Exact *P*-values are provided in the Source dataset.

### Limited biological roles for MpARF1 instability

Given the *in vivo* degradation of MpARF1, and the minor but significant effect of the R272Q mutation, a key question is whether MpARF1 instability is required for normal development or auxin response. Unlike effects of MpARF2 stabilization^21^, plants carrying *pro*Mp*ARF1*::Mp*ARF1*-*FL*^*R272Q*^-*mNG* did not show obvious deviations from normal growth (fig S2, F), suggesting a minor effect, if any. However, plants did contain a wild-type copy of Mp*ARF1*, complicating interpretation. We therefore replaced the endogenous MpARF1 with a R272Q mutant version, by introducing the mutant transgene into the Mp*arf1-4* loss-of-function mutant, which is insensitive to auxin treatment (fig. S3, A)^25^. While we found the R272Q mutation to cause slightly higher MpARF1 accumulation levels compared to WT in this background (Fig. 4, A to C; fig. S3, B), we did not observe substantial differences in phenotype when comparing Mp*arf1-4* complemented with WT or R272Q transgenes (fig. S3, C). Both complementation genotypes displayed similar restoration of auxin response (Fig. 4, D and E), and we did not find differences in sensitivity to auxin at a range of concentrations (fig. S4 A and B).

**Fig. 4.**
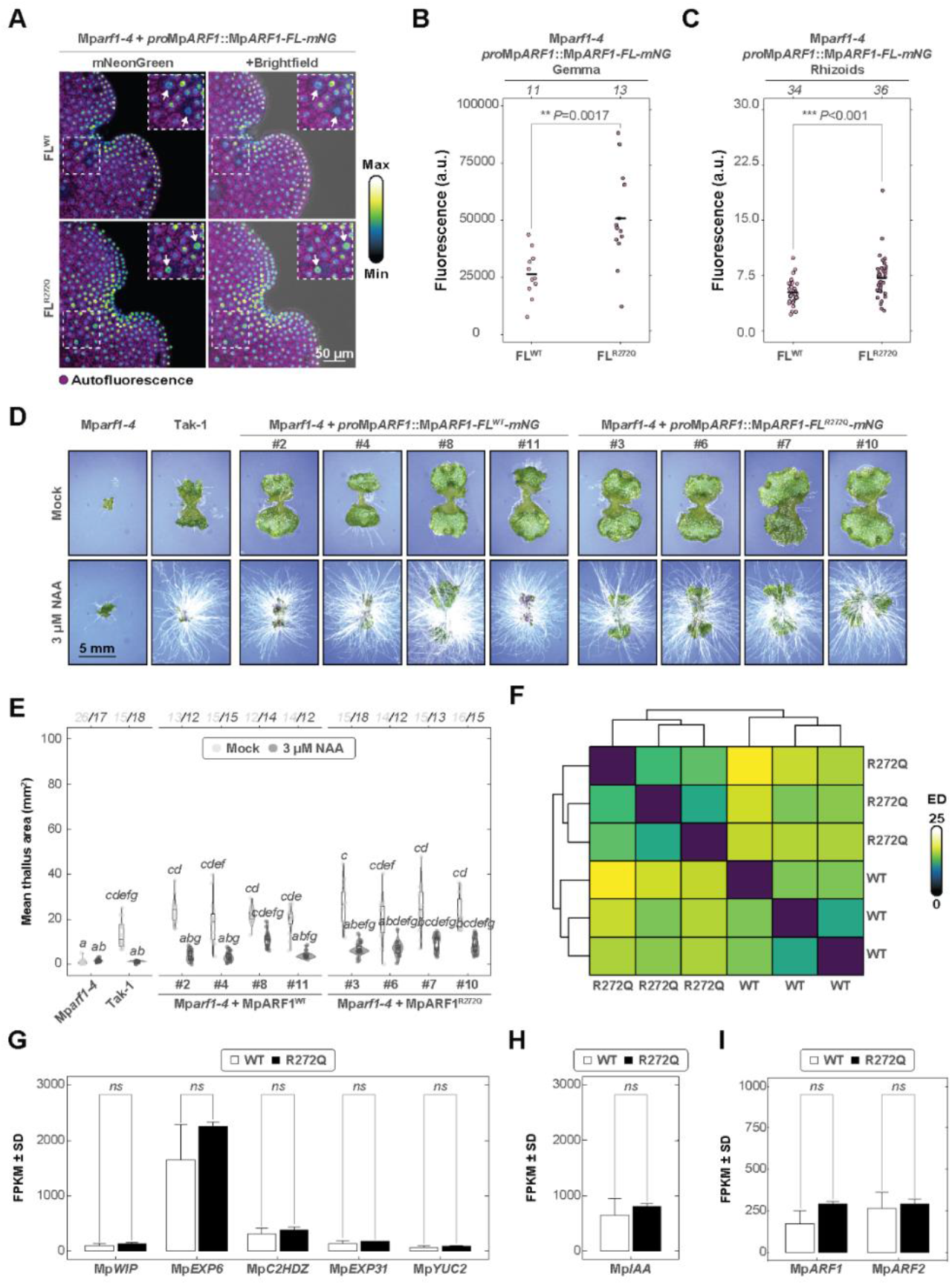
Functional analysis of MpARF1^R272Q^. (**A**) Representative confocal images of accumulation patterns of dormant gemmae expressing *pro*Mp*ARF1*::Mp*ARF1*-*FL*-*mNG* in a Mp*arf1-4* background. Magenta shows chloroplast autofluorescence. A portion of the image is outlined and enlarged, and nuclei of rhizoid initial cells are indicated with an arrow. (**B**) Quantification of overall fluorescence in several independent transgenic lines, of which one is shown in (A). Sample size is indicated above the figure and represents average fluorescence values for whole gemmae representing independent transgenic lines. (**C**) Quantification of nuclear fluorescence in rhizoid initial cells of plants shown in (A). Sample size is indicated above the figure, each replicate consists of a measurement from an individual nucleus of a rhizoid initial cell. (**D**) Qualitative assessment of auxin response in gemmalings grown for ten days on medium with Mock or NAA. Genotypes and independent transgenic lines are indicated above the figure. (**E**) Quantification of projected thallus area of the experiment shown in (D). Projected thallus area was compared per Kruskal-Wallis test (*H*(19)=243.77, *P*≈2.2×10^-16^) with a Dunn post hoc test (*P*<.05). Sample size is indicated above the figure and represents individual plants grown from gemmae. (**F**) Heatmap representing clustering of transcriptomes determined by RNA-seq of three independent biological replicates of Mp*arf1-4* mutant lines complimented with a *pro*Mp*ARF1*::Mp*ARF1*- *FL*^*WT*^-*mNG* or *pro*Mp*ARF1*::Mp*ARF1*-*FL*^*R272Q*^-*mNG* copy, respectively. ED; Euclidean distance. (**G**) Expression levels, represented as Fragments Per Kilobase Million of typical auxin-responsive genes in MpARF1^WT^ and MpARF1^R272Q^ complementation lines. (**H**) Expression levels, represented as Fragments Per Kilobase Million of Mp*IAA* in *pro*Mp*ARF1*::Mp*ARF1*-*FL*^*WT*^-*mNG* and *pro*Mp*ARF1*::Mp*ARF1*-*FL*^*R272Q*^-*mNG* complementation lines. (**I**) Expression levels, represented as Fragments Per Kilobase Million (FPKM) of Mp*ARF1* and Mp*ARF2* in MpARF1-FL^WT^ and MpARF1-FL^R272Q^ complementation lines. (B and C) Statistical differences were determined per T-test (*P<*.*05*) and sample size is indicated above the figures. (G to I) Bars represents means and error bars show standard deviation from the mean. Statistical differences were determined per T-test (*P<*.*05*).

To further examine the effects of altered protein accumulation on MpARF1 functions in gene expression regulation, we preformed RNA-seq analysis with *Mparf1-4* mutant, complemented either with a wild-type MpARF1 transgene, or with a transgene carrying the R272Q mutation. The two genotypes separated in hierarchical clustering (Fig. 4, F; fig. S5, A). We found 520 genes to be differentially expressed (172 up and 348 down), with no significant alterations in expression levels of known auxin-responsive genes (Fig. 4, F and G; fig. S5, B). Furthermore, no auxin-related GO terms were found to be enriched among differentially expressed genes (DEGs) (fig, S5, C), and no feedback regulation of Mp*ARF* or Mp*Aux/IAA* genes was detected (Fig. 4, H and I). These observations collectively suggest that reduction of MpARF1 degradation has a limited effect on MpARF1 function, both at the physiological and transcriptional levels.

### A developmental map of MpARF2 degradation

While MpARF degradation does not appear to be critically required for development, interfering with MpARF2 degradation dramatically alters thallus development^21^. This identifies MpARF2 degradation as a key control mechanism in auxin response. We next explored the extent to which this mechanism operates in controlling development throughout the Marchantia life cycle. To first map MpARF2 degradation without confounding effects of resulting plant phenotypes, we imaged plants expressing either *pro*Mp*ARF2*::Mp*ARF2*-*DBD*^*WT*^-*mNG* (DBD^WT^) or the *pro*Mp*ARF2*::Mp*ARF2*-*DBD*^*E297K+R300Q*^-*mNG* (DBD^E+R^) double mutant.

In early-stage developing dormant gemmae, the DBD^E+R^ mutant protein accumulated to high levels and broadly throughout the gemma, especially in the apical notch (Fig. 5, B). In later stage dormant gemmae, MpARF2 levels seem to be lower, but DBD^E+R^ seemed to be somewhat retained and is present in the apical notch and rhizoid initial cells (fig. S6, A and B). Intriguingly, we have previously shown that MpARF2 is usually absent or barely detectable in rhizoid initial cells^18^. The presence of DBD^E+R^ suggests therefore suggests that the Mp*ARF2* promoter is actually active in these cells but that MpARF2 abundance is regulated post-translationally through protein degradation. This is further supported by observation of signal in rhizoid initials in DBD^WT^ when treated with the proteasome inhibitor Bortezomib (Bz), as this treatment led to accumulation patterns similar to DBD^E+R^ (fig. S6, C). As MpARF2 expression is tightly linked to the meristematic apical notch zone^13,18^., we next decided to investigate MpARF2 degradation in apical notches of plants at a more precise temporal resolution. Whereas DBD^WT^ was not detected during the investigated stages, DBD^E+R^ was present throughout most cells (Fig. 5, C; fig. S6, D to E). We verified that this was due to proteasome-mediated degradation, and not a lack of expression, by treating plants with Bz, which resulted in accumulation (fig. S6, F). As stable lines expressing FL^E+R^ had problems in normal thallus growth, gemma- and gemma cup formation, we examined thallus-related structures such as air pores, air chambers, ventral scales and gemmae cups for DBD^WT^ degradation. DBD^E+R^ signal was observed widely throughout the thallus, mostly in photosynthetically active tissues, and less so in cells that lacked chloroplasts, and also in ventral scales (Fig. 5, D and E). DBD^E+R^ was present throughout various cell types in gemmae cups, including cells comprising the wings, the stalks and in cells which form the floor of the gemma cup (Fig. 5, D; fig. S6, G). In air chambers DBD^E+R^ accumulated to high levels in the air pore cells, in the filamentous photosynthetic cells located in the air chambers themselves, and sometimes in the surrounding ground tissue (Fig. 5, E).

**Fig. 5.**
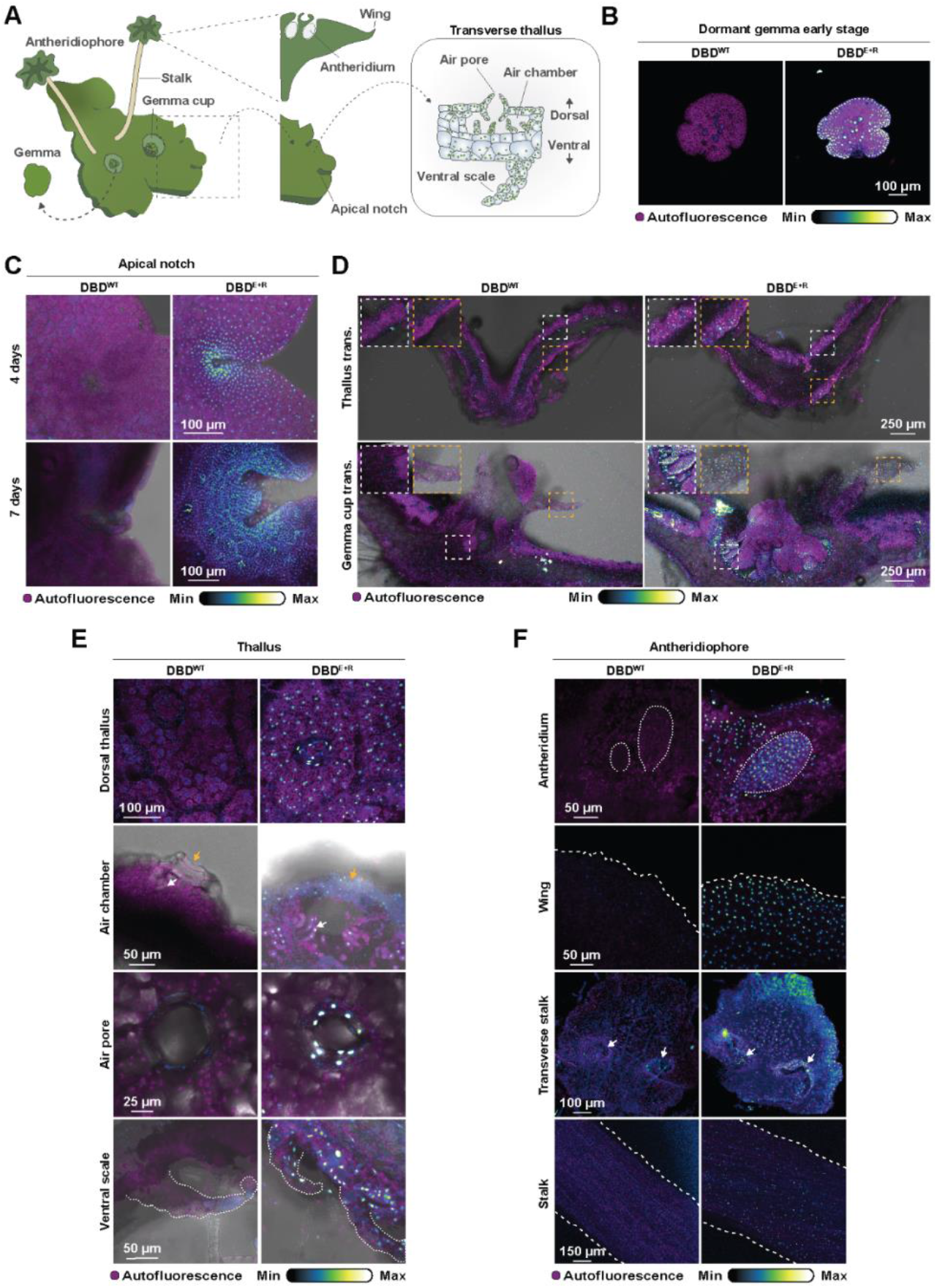
A spatial map of MpARF2 degradation. (**A**) Illustration of a mature *M. polymorpha* plant showing structures which were studied using confocal microscopy in this experiment. (**B to F**) Observations through confocal microscopy of accumulation patterns of a wild type (WT) and mutant (E297K+R300Q) copy of the MpARF2 DBD expressed from the Mp*ARF2* promoter fused to mNeonGreen, across different tissues. (**B**) Early stage gemmae. (**C**) Meristematic apical notches of plants grown from gemmae to a certain age, as indicated. (**D**) Transverse cross sections of a gemmae cup and a thallus respectively. Certain structures are enlarged in the white and yellow boxes. (**E**) Micrographs of mature thallus-related structures. Depicted here are a dorsal view of cells comprising the thallus, a dorsal view of an air pore, a transverse cross section of an air chamber (photosynthetic filaments (white arrow) and air pores (yellow arrow) are indicated), and ventral scales (outlined by white dotted lines). (**F**) Micrographs showing structures found in antheridiophores. Highlighted are antheridia (outlined in white dotted lines), the wing, a transverse cross section of the stem (invaginations containing rhizoid bundles indicated by a white arrow), and a lateral view of the stem.

Given that little is known about MpARF2 function and regulation in reproductive tissues, the gametangiophores, we investigated DBD^E+R^ accumulation in tissues which form the male antheridiophores (Fig. 5, A). DBD^E+R^ accumulated throughout the stem, mostly in peripheral cells of the stem and around the invaginations which harbor rhizoid bundles (Fig. 5, F). In cells which form the wing of the antheridiophore, DBD^E+R^ broadly accumulated (Fig. 5, F). DBD^E+R^ was especially enriched in cells which give rise to the jacket around newly developing immature antheridia (Fig. 5, F). Overall, these findings show that MpARF2 levels are tightly controlled through proteasomal degradation and that this degradation system is prominently active throughout the Marchantia lifecycle, enabling normal plant development.

### Extended functions of MpARF2 degradation

To study the relevance of MpARF2 degradation beyond initial gemma development, we developed a conditional version of the MpARF2-FL^E297K+R300Q^ protein expressed from its native promoter that can be activated, through excision of a floxed NLS-mScarlet3-tNOS cassette (abbreviated mARF2^flox^), by treatment with both Dexamethasone (Dex) and heat shock (HS). MpARF2 activation would be visible by loss of mScarlet3 fluorescence and gain of mNG fluorescence (Fig. 6, A). When mARF2^flox^ plants were exposed to HS alone, we observed some leaky editing in the absence of Dex; however, plants simultaneously treated with HS and Dex showed stronger and broader editing patterns (Fig. 6, B). Five days after induction, the leakiness of HS-treated plants seemed less severe, possibly due to selection against edited cells (fig. S7, A). The mARF2^flox^ plants did not display any phenotypic defects when compared to the Tak-1 wild-type background when not induced (fig. S7, B to E). To separate the effects of Dex and HS treatment, we included Tak-1 plants in all analyses, and observed that 1 hour of HS had minimal effects on development and thallus size (Fig. 6, C to D). We tested response to auxin in induced mARF2^flox^ gemmae and found clear auxin resistance and the characteristic phenotype found previously in stable MpARF2^E+R^ mutants (Fig. 6, C to D, fig. S7, F)^21^. This tells us that the mARF2^flox^ mimics the behavior of stable MpARF2^E+R^ mutants and that this system can be employed to study effects of MpARF2^E+R^ hyperaccumulation during thallus development.

**Fig. 6:**
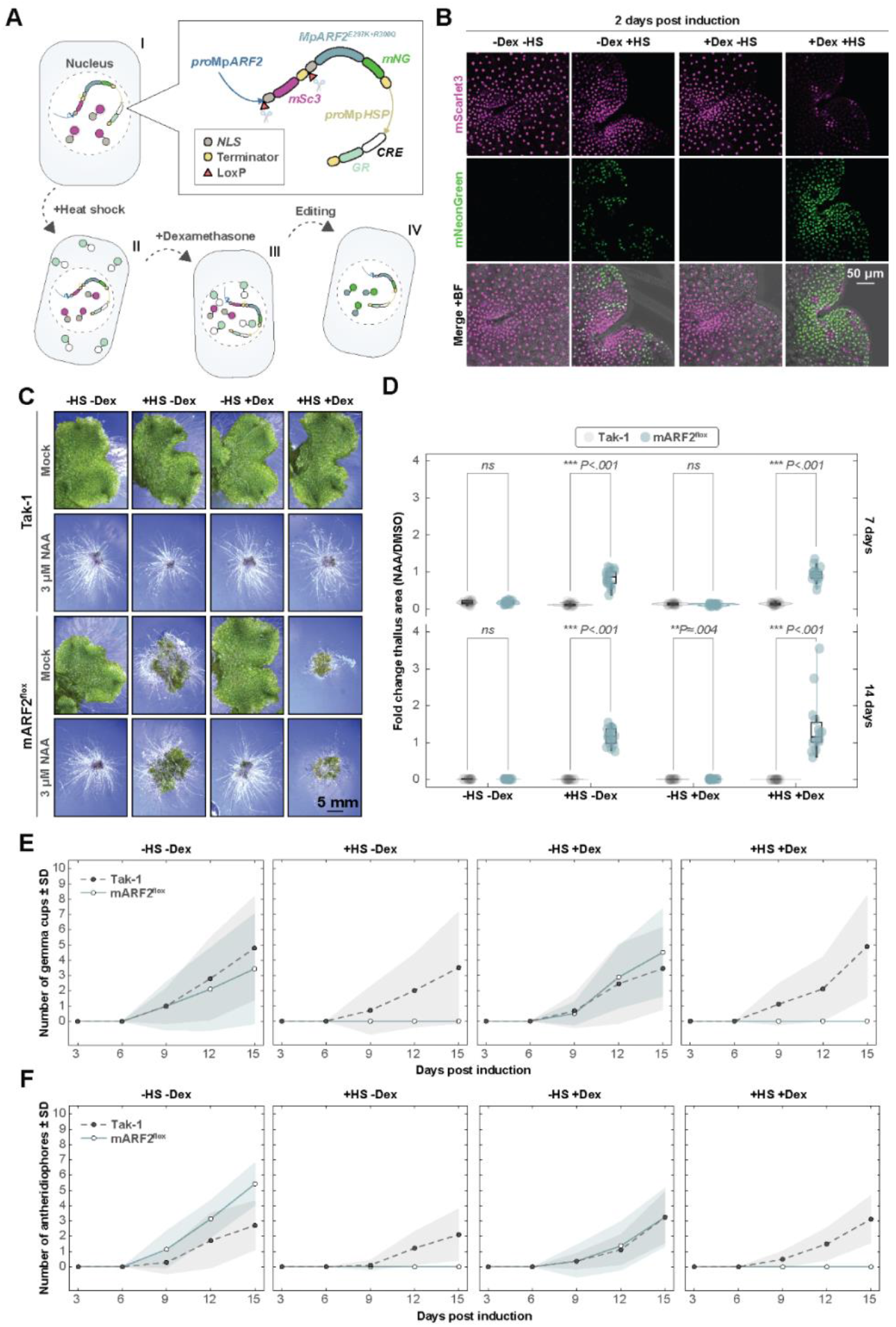
Conditional stabilization of MpARF2 reveals broad roles for its degradation in development. (**A**) Diagram illustrating the architecture of the inducible genetic construct, and how combined heat shock and Dexamethasone treatment leads to expression of mARF2^flox^. Abbreviations: mNG is mNeonGreen; mSc3 is mScarlet3; HSP is Heat Shock Protein; and GR is Glucocorticoid Receptor. (**B**) Observations of editing patterns of the inducible construct through confocal microscopy two days after induction with heat shock and Dexamethasone treatment. (**C**) Phenotype of Tak-1 and Inducible plants 14 days post induction with indicated treatments, grown on ½ Gamborg B5 medium supplemented with 3 μM NAA or Mock. (**D**) Quantification of projected thallus area of plants shown in (C) relative growth of NAA treated plants was calculated versus their Mock control (i.e. a low fold change means a strong reduction in size due to NAA treatment). Fold change in projected thallus size was compared per Mann-Whitney U test (*P*<.05) (**E**) Quantification of the number of gemma cups in Tak-1 and mARF2^flox^ plants measured during 15 days after induction with the indicated treatments. Shaded areas show standard deviations from the mean. (**F**) Quantification of the number of antheridiophores in Tak-1 and mARF2^flox^ plants measured during 15 days after induction with the indicated treatments. Shaded areas show standard deviations from the mean. (B to F) Abbreviations treatments: HS is heat shock; and Dex is Dexamethasone.

Auxin regulates gemma cup development^26^, a process which is difficult to study in stable mutants, as these generally do not form cups, or do so from abnormal thalli^21^. To study effects of MpARF2 stabilization on gemma cup development, we induced mARF2^flox^ after 7 days of cultivation and scored gemmae cup formation during the following 15 days. Strikingly, no gemmae cups were formed when mARF2^flox^ was induced even with only HS treatment (Fig. 6, E). Gemma cup formation was not impaired in the Mock or Dex treated control groups, and did not show morphological abnormalities (Fig. 6, E; fig. S8, A). In the previous experiments we found that MpARF2^E+R^ accumulates in antheridiophore associated tissues, whereas MpARF2^WT^ was maintained at low levels due to active degradation (Fig. 5, F). We therefore analyzed the effect of impaired MpARF2 degradation on antheridiophore formation using the mARF2^flox^ system. We grew plants for 14 days, and then induced both mARF2^flox^ and antheridiophore formation (with far red light), simultaneously. This developmental stage was also severely affected by induction of mARF2^flox^, as plants did not form normal antheridiophores when induced with HS or HS and Dex contrasting strongly with the Mock and Dex controls, which formed normal antheridiophores (Fig. 6, F; fig, S8, B). Some induced mARF2^flox^ plants, however, formed undefined outgrowths (fig. S8, B and C).

### MpARF2 degradation is required for biological function

Plants expressing a stabilized copy of MpARF2 are strongly affected in auxin response and development, a defect that persists throughout development. However, given that an endogenous copy is still present, it is unclear what the impact would be of preventing the only MpARF2 copy from being degraded. Earlier attempts to generate Mp*arf2* loss-of-function mutants failed to recover mutants, and we previously developed a floxed complementing MpARF2 to conditionally delete the only wild-type copy^13^. It is, however, impractical to complement this conditional mutant with specific protein versions. Recently, a stable Mp*arf2* allele was generated (Mp*arf2-1*, details will be published elsewhere), which now allows assessing the biological function of specific variants. To test the requirement for regulated MpARF2 degradation, we complemented the Mp*arf2-1* mutant either with an *pro*Mp*ARF2*::Mp*ARF2*-*FL*^*WT*^-*mNG* (MpARF2^WT^) or an *pro*Mp*ARF2*::Mp*ARF2*-*FL*^*E297K+R300Q*^-*mNG* (MpARF2^E+R^) copy. The uncomplemented Mp*arf2-1* mutant grows as a cell mass that retains the ability to proliferate and form rhizoid cells(Fig. 7, A). The mutant fails to produce a flat thallus and its associated structures such as gemmae cups and midribs (Fig. 7, A). Complementation with MpARF2^WT^ partially rescued the mutant phenotype (Fig. 7, A). Some of the transgenic lines obtained could form thalli with air pores, however, a full restoration of the phenotype was not observed, suggesting that some expression signals are likely missing from the transgenic construct. Obtained lines still exhibited slower growth, abnormal apical notch formation and branching patterns when compared to Tak-1 (Fig. 7, A). Some lines were able to produce gemmae cups and gemmae, one line produced gemmae without clearly defined notches, resulting in a round morphology (Fig. 7, B; fig. S9, A). These plants exhibited a relatively normal response to auxin (Fig. S9, B). In contrast, MpARF2^E+R^ plants failed to produce thalli with gemmae cups, and still appeared more or less like the Mp*arf2-1* mutant, even though they were able to grow faster (Fig. 7, A). Excessive accumulation of MpARF2 leads to repression of the auxin response. Either way, these results show that regulated degradation is required for MpARF2 function *in vivo*.

**Fig. 7.**
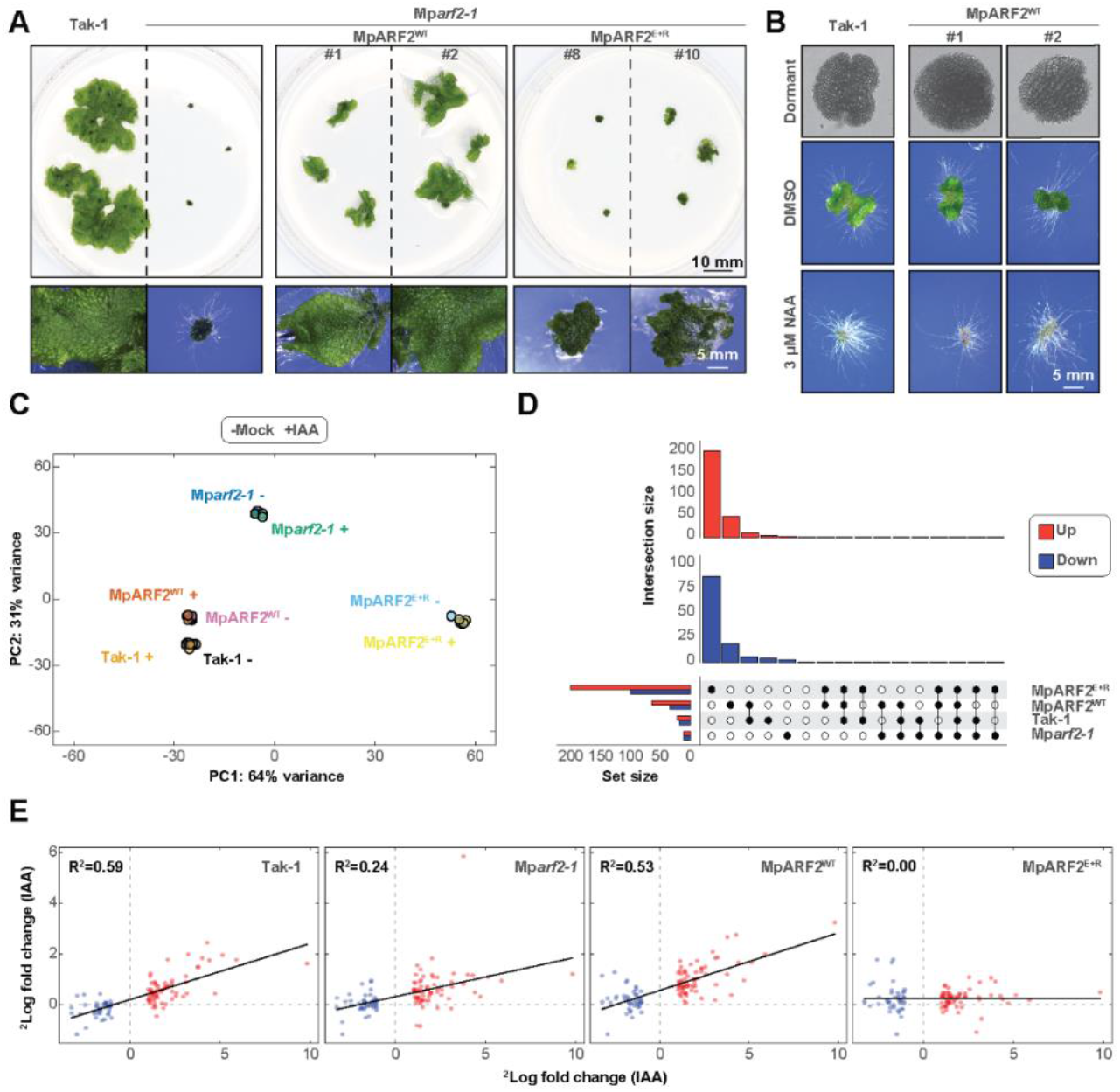
MpARF2 proteolysis is required for biological function. (**A**) Phenotypes of 20-day-old plants of the indicated genotypes, grown from tissue fragments. The lower panels show an enlargement of the plants imaged above. (**B**) Gemma morphology and qualitative analysis of auxin response of ten-day-old gemmalings of Mp*arf2-1* mutant lines complemented with MpARF2^WT^ grown on NAA or Mock. (**C**) Principal component analysis following transcriptome analysis of Tak-1, Mp*arf2- 1*, and Mp*arf2-1* complementation lines (MpARF2^WT^ and MpARF2^E297K+R300Q^, respectively) treated with IAA (indicated with +) or Mock (indicated with −). (**D**) Upset plot showing unique and overlapping differentially expressed genes (−0.5<^2^Log fold change>0.5, *P*_adj_<.05) in the IAA-treated group versus the Mock control, per genotype. (**E**) Relationship between fold changes of a set of auxin-responsive genes identified by Ref.^12^ on the x-axis, and fold changes found in this experiment on the y-axis. R^2^ value depicts fit of the linear regression model.

To address the impact of reduced MpARF2 degradation on MpARF2 function in transcriptional regulation, we grew fragments of Tak-1, Mp*arf2-1*, MpARF2^E+R^, and MpARF2^WT^ plants for 14 days, depleted endogenous auxin with the synthesis inhibitors Yucasin and L-Kynurenin^27^, followed by treatment with 1 μM Indole-3-Acetic acid (IAA) or Mock for one hour. RNA-sequencing results showed strong clustering by genotype, eclipsing effects of the IAA treatment (Fig. 7, C; fig. S10, A). Notably, MpARF2^WT^ clustered together with Tak-1, whereas MpARF2^E+R^ clustered separately (Fig. 7, C). MpARF2^E+R^ had the highest number of DEGs, showing no overlap with the other genotypes, indicating that these DEGs are not genes that are auxin-responsive (Fig. 7, D). This was confirmed through GO-analysis, which showed that no auxin-related terms were found among terms enriched in genes differentially expressed in MpARF2^E+R^ (fig. S10, B). We analyzed the expression of a set of previously described IAA-responsive DEGs^12^ and found that both Tak-1 and MpARF2^WT^ showed highly similar expression of these genes (Fig. 7, E). Unexpectedly, Mp*arf2-1* did not show a hypersensitive response, but rather a slightly dampened response to auxin (Fig. 7, E). The transcriptional response to IAA was completely abolished in MpARF2^E+R^, confirming that the physiological effects observed in these mutants are due to both a very strong repression of the auxin response system as well as unlicensed regulation of auxin-unrelated genes (Fig. 7, E). These results confirm that MpARF2 degradation is essential for proper transcriptional homeostasis and responses to auxin, which, combined with effects on physiology, confirm its importance.

## Discussion

Auxin is an important signaling molecule, deeply intertwined with plant development, and largely relies on transcriptional reprogramming. ARF proteins are an essential component of auxin signaling and their activity must be tightly regulated. Here, we report on significant divergence in degradation dynamics and functional consequences of MpARF family members, thereby offering new insight into post-translational tuning of auxin signaling. More specifically, we demonstrate that a conserved arginine residue in the DBD is required for MpARF proteolysis. Our results further indicated that, although the A-class MpARF1 is subjected to proteolysis, interference with its degradation through this arginine only resulted in mildly increased protein levels with negligible consequences on MpARF1 function and consequently, auxin signaling and plant development. This is in strong contrast to our finding that B-class MpARF2 degradation is critically required for biological function. We show that MpARF2 accumulates in essentially all cell- and tissue types across the Marchantia lifecycle when degradation is impaired, leading to severely derailed developmental programs. Finally, we demonstrate that MpARF2 function is contingent on its degradation through the proteosome, as dysregulation through proteolysis results in uncontrolled activity and a completely impaired transcriptional auxin response. Our findings point toward degradation mechanisms whose specificity are a product evolutionary adaptation in the embryophyte lineage.

The A- and B-class ARFs represent two clades with deep evolutionary roots, and their common ancestry dates back to the very early stages of colonization of land by plants^23^. Most likely, the duplication of an ancestral AB-class ARF, which took place close to the origin of land plants (an event which likely occurred in ancestor of all embryophytes), resulted in the lineages giving rise to the respective A- and B-class clades^23,28^. The A- and B-class ARFs then acquired specialized, antagonizing functions, wherein A-ARFs gained the ability to act as transcriptional activators while B-class ARFs tend to function as repressors^13,14,23^. In earlier work, we hypothesized that the DBD-mediated ARF instability arose in the AB-ARF ancestor of the A- and B-ARF clades, roughly 500-600 MYA^21^. Here we show that the same motif identified in B-ARF MpARF2 has similar, instability-conferring, properties within the context of the A-ARF MpARF1, thus providing further evidence supporting this hypothesis. However, the results presented here indicate that, although the DBDs of A- and B-ARFs are structurally and functionally deeply conserved, proteasomal degradation through signals embedded in this DBD have either gained importance in B-ARFs or lost importance in A-ARFs^11,12^. Possibly, the ubiquitin-ligase adapter protein which targets ARFs has coevolved with B-ARFs, resulting in higher binding affinity, and thus increased susceptibility for degradation. There is also the possibility of additional cofactors, which regulate substrate accessibility by the ubiquitin-ligase, which have evolved to target B-ARFs but not, or to lesser extent, A-ARFs. Such innovations would then enable more precise spatiotemporal control over MpARF2 levels, thereby permitting evolution of stronger repression capabilities and/or the ability to regulate new gene sets, whereas MpARF1 would only be weakly targeted and therefore evolved weaker abilities. Alternatively, B-ARFs might undergo post translational modifications which decrease its stability, e.g. by increasing affinity for the ubiquitin-ligase, whereas A-ARFs might be modified such that they are more protected from degradation, these theories require testing at the level proteomics level. Many of these questions would be answered upon identification and proper functional characterization of said ubiquitin-ligase, which remains unidentified. A previously identified ubiquitin-ligase AUXIN RESPONSE FACTOR F-BOX1 (AFF1), which targets the Arabidopsis A-class ARFs AtARF7 and AtARF19^19^, is not conserved in Marchantia and therefore, another ubiquitin-ligase must be involved.

Surprisingly, hampered MpARF1 degradation had barely any noticeable effects on MpARF1 function and plant development. We also observed that full-length MpARF1 proteins expressed from the *pro*Mp*UBE2* promoter accumulated to lower levels than each domain separately. Potentially, other domains of MpARF1 (the MR and PB1 domain) could, through a potential joint effort, contribute to its proteostasis. There are examples of A-class ARFs relying on their MR or PB1 domains for degradation^29,30^, so possibly, A-class ARFs have evolved multiple ways to be routed to the proteasome which are lacking or less-strong in B-class ARFs. Furthermore, A-class ARF activity can also be inhibited through interactions with the Aux/IAA co-repressors, providing a reason for the lack of physiological effects when MpARF1 accumulation increased. Why then, did MpARF1 not lose degradation through the DBD? Since evolution of the DBD is known to be under high levels of evolutionary constraint, with little room for sub- or neofunctionalization^12^, it is likely that the motif responsible for degradation is required for structural integrity of the MpARF1 DBD, and DNA-binding, thus preventing changes to occur.

In summary, the work presented here reveals that, despite deep conservation of the ARF DBDs, the role of proteasome-mediated degradation has strongly diverged between A- and B-class ARFs, which likely reflects distinct evolutionary pressures and regulatory demands. The strict functional dependency of MpARF2 on proteolysis, which contrasts with the dispensability of MpARF1 degradation, may have been enabled or enforced by co-evolution with the degradation machinery or acquisition of unique regulatory mechanisms between the two classes during land plant evolution. Further deciphering of the precise mechanism of ARFs by the ubiquitin-proteasome system, which includes identifying the responsible ubiquitin-ligase, remains a crucial hurdle to address remaining questions, as answering these questions will be essential to fully understand tuning of auxin signaling. Ultimately, our findings suggest that diversification of functional requirements of ARF proteolysis may help explain how the simple signaling molecule auxin can orchestrate such a wide range of developmental processes in a precisely controlled manner.

## Materials and Methods

### Protein structure prediction & visualisation

The MpARF2 crystal structure was retrieved from PDB (https://www.rcsb.org/structure/6SDG)^13^. To model the MpARF1 structures, as well as the swap and mutated versions of the MpARF2 DBD, we employed AlphaFold2-multimer v2.2.2 with the default settings^31^. Structural visualisation was performed using Pymol (v2.3.4) software.

### Plasmid construction

Plasmids and oligonucleotides which were generated for this study can be found in Table S1 and S2, respectively. Promoters and cDNA-derived sequences were amplified through PCR from in-house plasmids mainly from Ref.^13,21,32^. The pMpGWB100 and pMpGWB300 backbones for the plasmids were from Ref.^33^. The motif swap of MpARF1 R272Q into the MpARF2 DBD-mNeonGreen fusion was generated based on the alignment in Ref.^21^.

To generate the plasmids expressing MpARF2 fusion proteins (the mutagenesis screen, motif swaps and Mp*arf2-1* complementation) we used a *pro*Mp*ARF2* pMpGWB100 backbone (which contains an ~1.9kb *pro*Mp*ARF2* promoter upstream of an XbaI site, an in-frame mNeonGreen (mNG) coding sequence, followed by a tNOS^21^. Mutations were generated through site-directed PCR.

For MpARF1 fusion proteins, we generated *pro*Mp*UBE2* pMpGWB100 plasmids by inserting the ~3kb Mp*UBE2* promoter^32^ into the XbaI site of pMpGWB100 together with the respective MpARF1 domains fused to mNG. For the *pro*Mp*MpARF2*::*NLS-*Mp*ARF1-DBD*^*WT*^*-mNG* and *pro*Mp*MpARF2*::*NLS-* Mp*ARF1-DBD*^*R272Q*^*-mNG* pMpGWB100 plamids we used the *pro*Mp*ARF2* pMpGWB100 backbone described above, and inserted the MpARF1 domains into the XbaI site. For MpARF1 complementation assays, we first cloned the full length MpARF1 sequences, C-terminally fused to mNG, into pDONR221 to generate pENTR221 MpARF1-FL^WT^-mNG and pENTR221 MpARF1-FL^R272Q^-mNG. Next, expression vectors were generated through LR-reaction of entries and pHKDW046 (housed in the pMpGWB300 plasmid), which contains a ~4900bp Mp*ARF1* promoter upstream of the attR sites^13,33^.

To clone mARF2^flox^, we synthesized a fragment encoding a floxed SV40 Nuclear Localization Signal (NLS) fused to mScarlet3 (mSC3) and a tNOS terminator (loxP-NLS-mSc3-tNOS-loxP; Fig. 6, A). We used our XbaI-digested *pro*Mp*ARF2* pMpGWB100 backbone (which contains an ~1.9kb *pro*Mp*ARF2* promoter cloned into the XbaI site, with the XbaI site regenerated) to insert the (from 5’ to 3’): loxP-NLS-mSc3-tNOS-loxP fragment downstream of *pro*Mp*ARF2* together with the *MpARF2*-*mNG*-tNOS CDS and *pro*Mp*HSP*::*Cre*-*GR-*tNOS amplified from Ref.^34^.

### Plant growth conditions and transformation

For this study *Marchantia polymorpha* Takaragaike-1 (Tak-1) was used as the wild-type. Plants were grown on ½ strength Gamborg B5 medium (1% agar) at 22°C with 40 μmol photons m^−2^s^−1^ continuous white fluorescent light. Tak-1 was transformed using agrobacterium-mediated delivery of transgenic constructs as described by Ref.^35^. Transgenic plants were selected on ½ Gamborg B5 (1% agar) with 10 mg/L hygromycin and 100 μg/ml cefotaxime or 0.5 μM chlorsulfuron.

### Analysis of genetically induced expression of MpARF2^E+R^

Plants at the indicated developmental stages were placed in ½ strength Gamborg B5 medium, and treated with a 1 hour 37°C heat shock (HS) and/or 10 μM Dexamethasone (Dex, Sigma-Aldrich) dissolved in ethanol^34^. Afterwards, plants were placed back in their normal growth condition to recuperate and phenotype parameters were scored. For determination of auxin response, gemmae were placed onto their respective media and immediately treated with 1 hour of HS.

For scoring of gemma cup formation, plants were grown from gemmae on ½ strength Gamborg B5 medium plates (1% agar) containing Mock (Ethanol) or Dex (10 μM) for seven days and were subsequently induced with 1 hour of HS. The number of gemma cups per plant were counted every three days using a Leica M205 stereomicroscope.

For scoring of antheridophore emergence, plants were grown from gemmae for 14 days grown from gemmae in plastic boxes, with ventilation, on ½ strength Gamborg B5 medium (1% agar) containing Mock (Ethanol) or Dex (10 μM). Plants were then induced with 1 hour of HS, and subsequently placed in 22°C with 40 μmol photons m^−2^s^−1^ continuous white fluorescent light supplemented with far-red light (50% power). The number of antheridiophores per plant were counted every three days using a Leica M205 stereomicroscope.

### RNA-sequencing & Transcriptome analysis

For the Mp*arf2-1* mutant complementation, the indicated genotypes were ground into small fragments using a tissue homogenizer (Fisherbrand^TM^ 150 homogenizer) and plated on ½ strength Gamborg B5 medium (1% agar) and grown at 22°C with 40 μmol photons m^−2^s^−1^ continuous white fluorescent light for 14 days. Regenerated plant fragments were then cultured for 6 hours in liquid ½ strength Gamborg B5 medium with 50 μM Yuc and L-Kyn, respectively. Then, plantlets were washed and cultured in liquid ½ strength Gamborg B5 with 1 μM Indole-3-Acetic acid (Alfa Aesar), or an equal volume of DMSO in the mock condition, for 1 hour. Afterwards, plants were harvested, flash-frozen in liquid nitrogen and subsequently ground into a fine powder.

For Mp*arf1-4* mutant complementation analysis plants were grown for eight days from gemmae on standard ½ strength Gamborg B5 medium, without any treatments. Plants were then harvested and directly frozen in liquid nitrogen and processed as described below.

For RNA extraction, ground tissue was incubated in 1 mL of TRIzol reagent for 5 min. at Room Temperature (RT). Next, 200 μL chloroform was added, samples were vortexed and incubated for an additional 3 min. at RT. Following this, samples were centrifuged for 15 min. at 12000g and 4°C, and the upper aqueous phase containing the RNA was transferred to a new tube and precipitated with 500 μL isopropanol for 10 min. at RT. Samples were then transferred to pink columns and processed further using the Qiagen RNeasy RNA purification kit, according to the manufacturer’s instructions. RNA was sequenced using Illumina NovaSeq X plus by Novogene and checked for quality using FastQC. Reads were then mapped to the *M. polymorpha* genome (v7.1) using Hisat2 and read counting was performed using FeatureCounts. Read count normalization and differential expression analysis was performed through DEseq2, in which we excluded genes with a cumulative read count of <30 reads across all samples. Gene Ontology analysis was achieved using PlantRegMap using genelists as input of up- and downregulated genes respectively. In the Mp*arf1-4* mutant complementation experiment, Differentially Expressed Genes (DEGs) were defined as having a ^2^Log Fold Change below and above −1 and 1, respectively (*P*_*adj*_<.05). For the Mp*arf2-1* mutant complementation and auxin response experiment, we defined a ^2^Log Fold Change below and above −0.5 and 0.5 as cutoffs, respectively (*P*_*adj*_<.05).

### Confocal microscopy & Assessment of fusion protein stability

Confocal microscopy was carried out with a Leica SP8X-SMD confocal microscope fitted with hybrid detectors and a 40 MHz pulsed white-light laser. Data acquisition was performed using LasX (v3.5.7.23225) software. The fluorophore mNeonGreen was excited using 488 nm laser line, and mScarlet3 with a 561 nm laser line. Fluorescence was detected between 500-570 nm (mNeonGreen) or 573-640 nm (mScarlet3) with hybrid detectors set to photon counting mode with 1.00-24.50 ns time-gating active to suppress background fluorescence. Z-stacks were acquired using an HC PL APO 20x/0.75 water immersion or APO CS 10x/0.40 air objective. ImageJ (v1.52) was used to generate maximum-intensity projections, measure fluorescence intensities and produce micrographs for figures.

When investigating stability of MpARF2-DBD-mNG fusion proteins carrying different mutations, we screened T1 transgenic lines for fluorescence by imaging the apical notch region of dormant gemmae. When no fluorescence was detected, we treated these gemmae with Bortezomib (Cayman Chemical) for 24 h or the solvent DMSO (Sigma-Aldrich) as a Mock control, and checked for fluorescence again. Transgenic lines which showed no nuclear mNeonGreen fluorescence after Bortezomib treatment were excluded from the analysis. To quantify average fluorescence, we generated ROIs by tracing the outline of gemmae. To measure the background signal we created ROIs which excluded nuclei, obtained the average intensity and corrected the original measurement by subtraction of the average background signal per pixel. For quantification of rhizoid initial cells we drew ROIs around the respective nuclei and measured the average intensity.

When imaging gemmae over time, an imaging chamber was created by placing five 25 μL Gene Frames (Thermo Scientific) onto an objective glass, which was then filled with ½ strength Gamborg B5 medium (1% agar) with the indicated treatments. Gemmae were placed onto the medium, and the imaging chambers were filled with Perfluordecalin (Sigma-Aldrich) to allow gas exchange. Chambers were sealed with a coverslip, when not being imaged, slides were placed in their normal growth conditions.

### Phenotype analysis

For experiments assessing fluorescence accumulation, non-chimeric transgenic lines were established by growing a G_1_ generation from gemmae of independent plants that appeared resistant to initial selection on hygromycin or chlorsulfuron (T_1_ generation). Gemmae from the G_1_ generation were used for experiments. To assess the phenotypes of plants expressing the full length MpARF1 fusion proteins, we reported on phenotypes of 20-day old plants. For growth assays, dormant gemmae were micro dissected from the gemmae cups/thallus and lightly placed onto the respective medium. For general overviews or thallus size quantifications, pictures were taken with a Canon EOS250D camera and thallus area was measured using ImageJ (v1.52). For higher magnification images a Leica M205 stereomicroscope was used.

### Statistics & reproducibility

Statistical tests were performed in R (v4.2.1). Tests, parameters, additional statistical information and the nature of biological replicates are provided in figure descriptions and Source Data. Sample sizes are shown within the figures themselves and their nature is described in figure descriptions. No statistical method was used to predetermine sample size, sample sizes were determined according to field standards. Experiments were not randomized. Representative micrographs, are shown which reflect the average values in the corresponding quantitative experiments (as indicated).

## Acknowledgments

We would like to thank Jorge Hernández-García, Jiashu Chu, and Juriaan Rienstra for helpful discussions regarding this work. Furthermore, we to thank Danilo dos Santos Pereira and Jasper Lamers for assistance with the transcriptome analysis, and Emi Hainiwa, Hidemasa Suzuki, and Takayuki Kohchi for their efforts in isolating the Mp*arf2-1* mutant.

## Author contributions

Conceptualization: MDR, DW

Methodology: MDR, DW

Investigation: MDR, EH, DW, RN

Visualization: MDR

Supervision: JWB, DW

Writing—original draft: MDR, DW

Writing—review & editing: MDR, DW

## Data and materials availability

Plasmids generated and used in this study are available upon request from the corresponding author. As long-term storage of *Marchantia polymorpha* lines is problematic, we can provide some lines analyzed in this study, contact the corresponding author for inquiries. We provide a source data file containing the raw data and statistical output for the quantitative experiments contained in this paper. Raw RNA-sequencing data is deposited on NCBI and can be accessed through BioProject accession PRJNA1313408.

