## Supplementary Material for "Diversification of functional requirements for proteolysis of Auxin Response Factors"

Martijn de Roij *et al.*

**This PDF file includes:**

Figs. S1 to S10
Tables S1 and S2

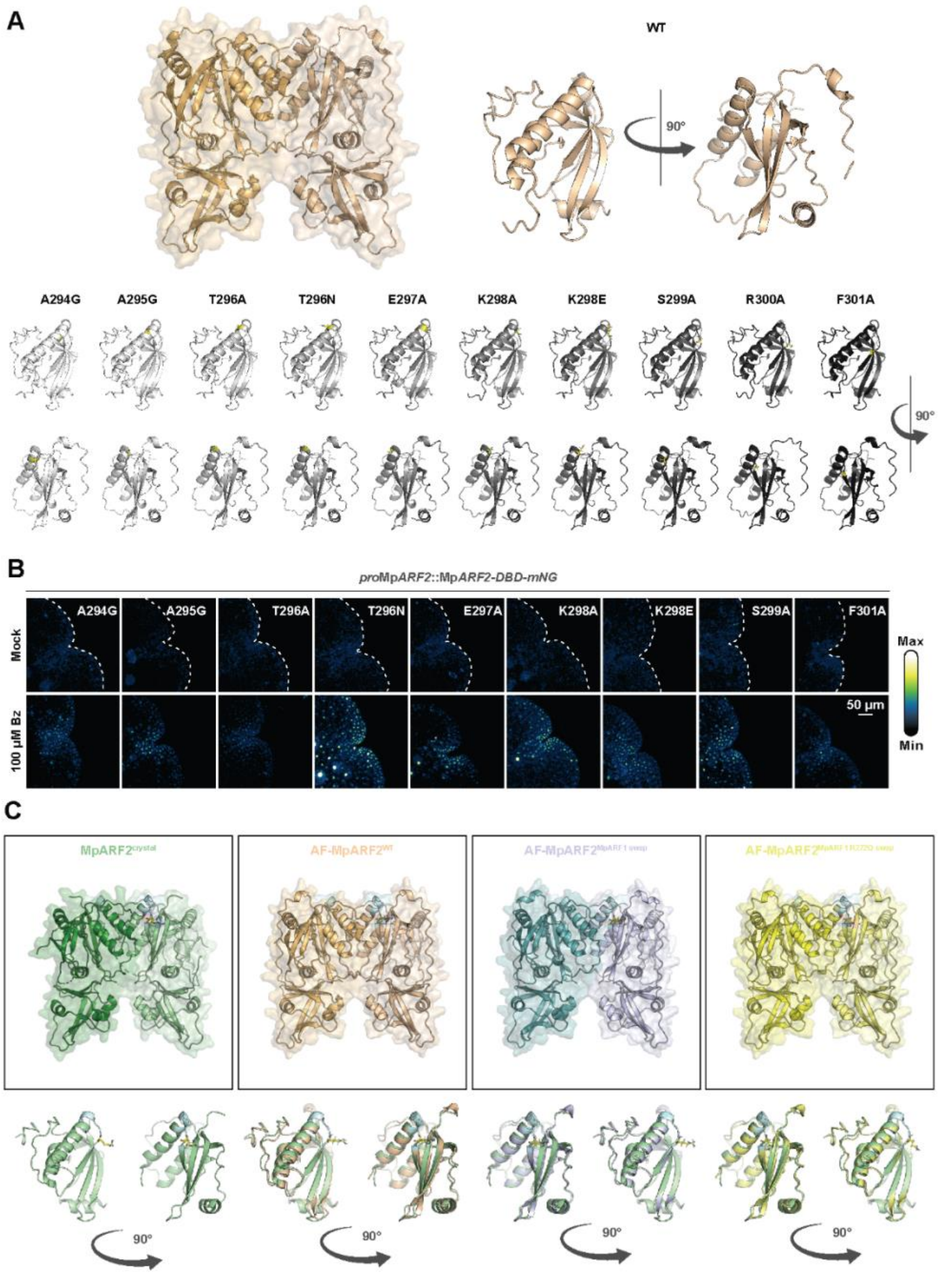


**Fig. S1. Structural prediction and proteasomal degradation of MpARF2 variants.**

(**A**) Structural models generated by AlphaFold2 of the MpARF2 DBD (orange) as a homodimer, and the structure containing the loop with relevant residues is outlined and enlarged. Below are AlphaFold2 models of the MpARF2 DBD with indicated point mutations, the relevant residue is colored by element (yellow). (**B**) Accumulation patterns of *pro*Mp*ARF2*::Mp*ARF2-DBD-mNeonGreen* fusions, carrying the indicated mutations, treated for 24 hours with Bortezomib (Bz) or Mock. (**C**) Structure of the MpARF2 DBD crystal (green) separately and overlain on AlphaFold2 models of a WT (orange) and mutant versions of the MpARF2 DBD as indicated.

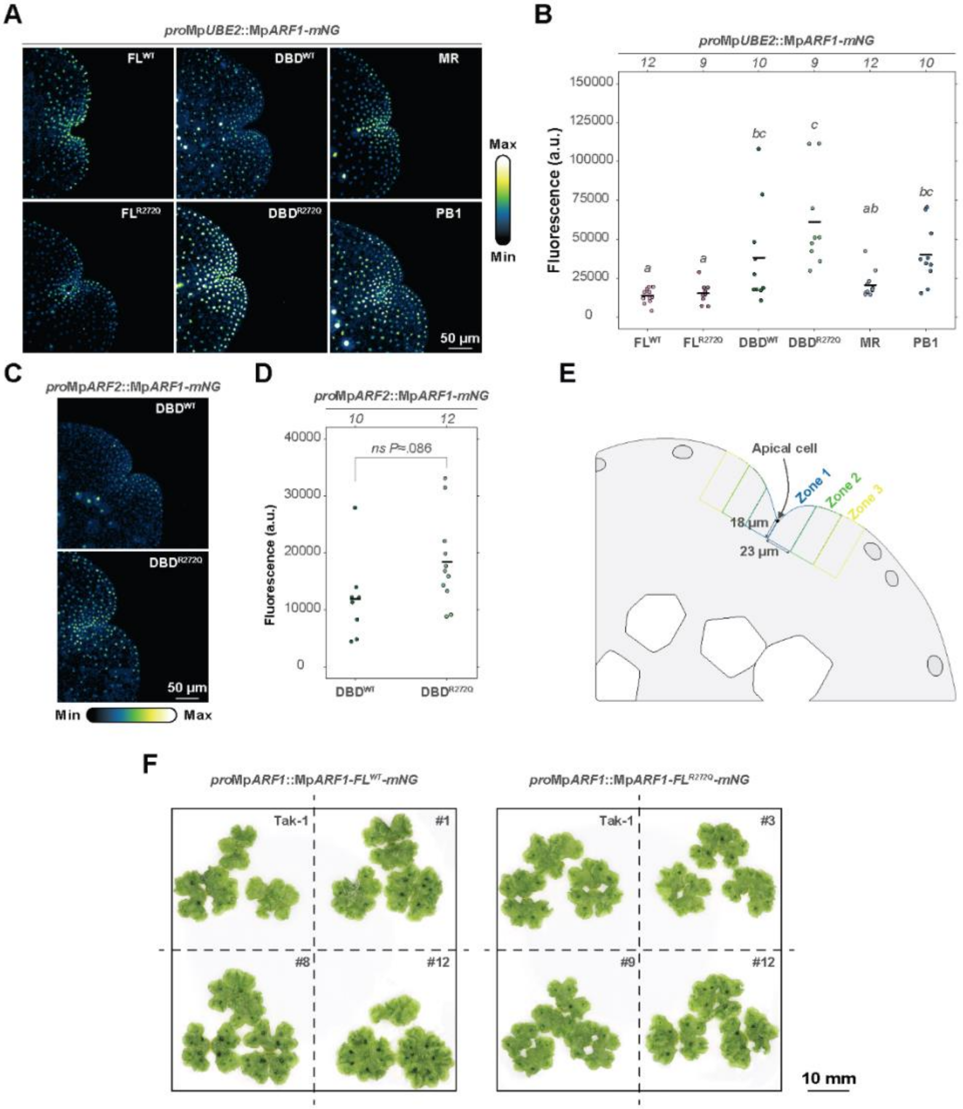


**Fig. S2. Determinants of MpARF1 degradation, and impact of mutation on phenotype.**

(**A**) Representative confocal images of accumulation patterns of dormant gemmae expressing protein fusions of MpARF1 to mNeonGreen as indicated. Fusion proteins are expressed from the constitutive Mp*UBE2* promoter. (**B**) Quantification of overall fluorescence in a number of independent transgenic lines of which one is shown in (A). Sample size is shown above the figure and corresponds to measurements of a gemma each from an independent transgenic line. Letters in italics indicate significant differences determined per one-way ANOVA ($F$=(5,56)=12.83, $P$≈2,54x10$^{-8}$) and Tukey HSD post hoc test with Bonferroni correction ($P$<.05). (**C**) Representative confocal images of MpARF1 DBD-mNeonGreen fusions expressed from an *pro*Mp*ARF2* promoter. (**D**) Quantification of overall fluorescence in gemmae, same experiment as (C), compared by T-test ($P$<.05). Sample size is shown above the figure and corresponds to measurements of a gemma each from an independent transgenic line. (**E**) **I**llustration showing an arbitrary subdivision of the apical notch meristematic region into three zones, positioned relative to the apical cell. (**F**) Representation of the overall morphology of 20 day old Tak-1 plants and Tak-1 plants carrying an extra copy of *pro*Mp*ARF1*::Mp*ARF1-FL*$^{WT}$*-mNG* or *pro*Mp*ARF1*::Mp*ARF1-FL*$^{R272Q}$*-mNG*. (B and D) Exact $P$-values are provided in the Source dataset. (A and B) Abbreviations of MpARF1 domains: FL; Full Length, DBD; DNA-binding domain, MR; Middle Region, PB1; Phox and Bem 1.

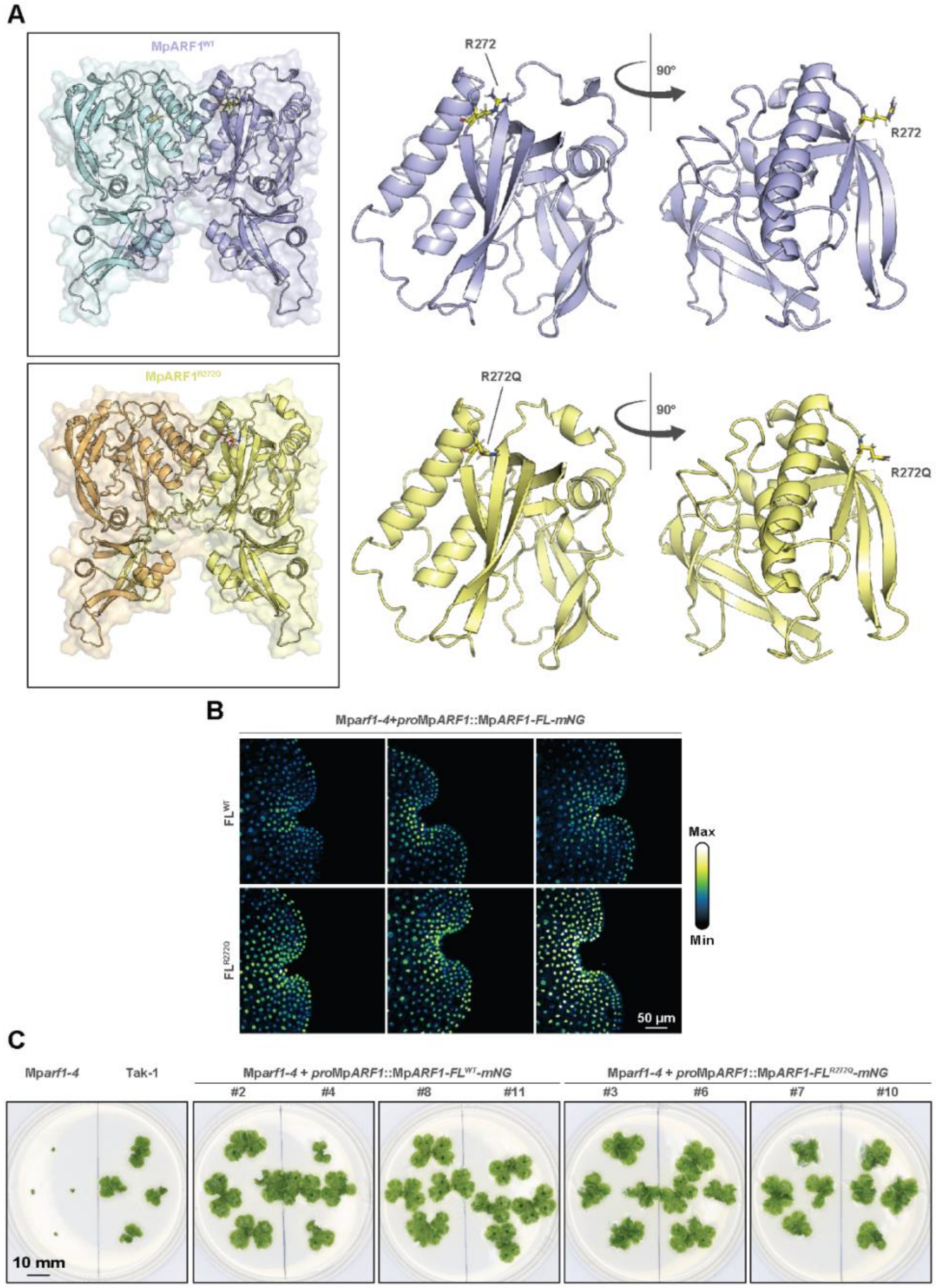


**Fig. S3. Structural modeling, protein accumulation and biological activity of MpARF1$^{R272Q}$ mutation.**

(**A**) AlphaFold2 models of the MpARF1 DBD structure as a homodimer (WT in blue and purple, R272Q mutant in orange and yellow), and a zoomed in region harboring the R272 or R272Q residue (highlighted). (**B**) Three additional, independent transgenic lines of the *Mparf1-4* mutant complimented with *pro*Mp*ARF1*::Mp*ARF1-FL$^{WT}$-mNG* (FL$^{WT}$) and *pro*Mp*ARF1*::Mp*ARF1-FL$^{R272Q}$-mNG* (FL$^{R272Q}$) proteins, to supplement micrographs shown in Fig. 4, A. These lines represent the means of Fig. 4, B. (**C**) Overall phenotypes of plants aged 20 days, grown from gemmae, of the indicated phenotypes.

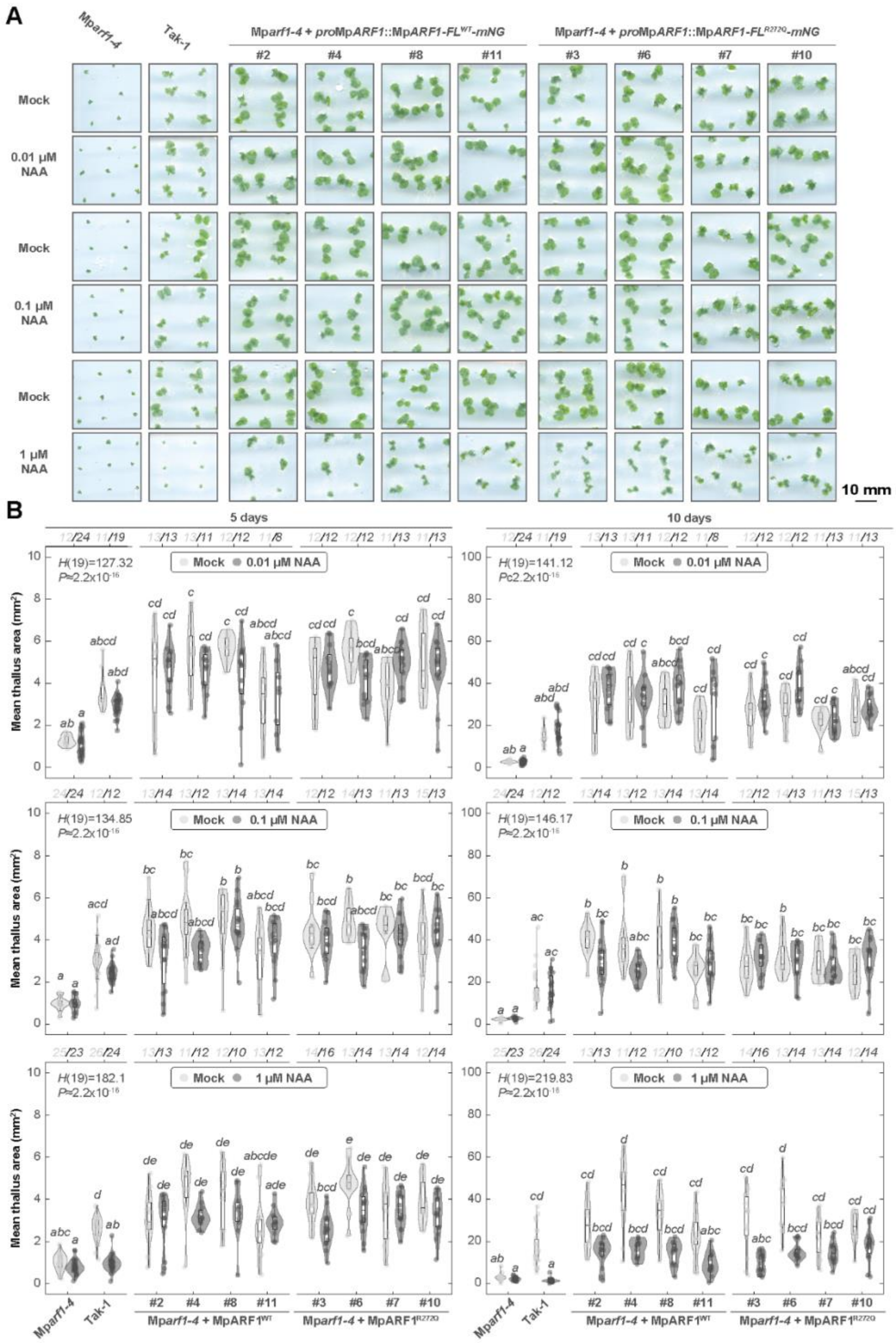

**Fig. S4. Quantification of auxin response in the MpARF1$^{R272Q}$ mutant.**

(**A**) Overview of general growth of the indicated genotypes (top) grown on a range of NAA concentrations (left) for five days. (**B**) Quantification of projected thallus area of plants of indicated genotypes grown on medium with 1, 0.1 and 0.01 µM NAA for five and seven days, respectively. Projected thallus areas was statistically compared by Kruskal-Wallis with Dunn post hoc test ($P$<.05), test statistics are shown within the figures.

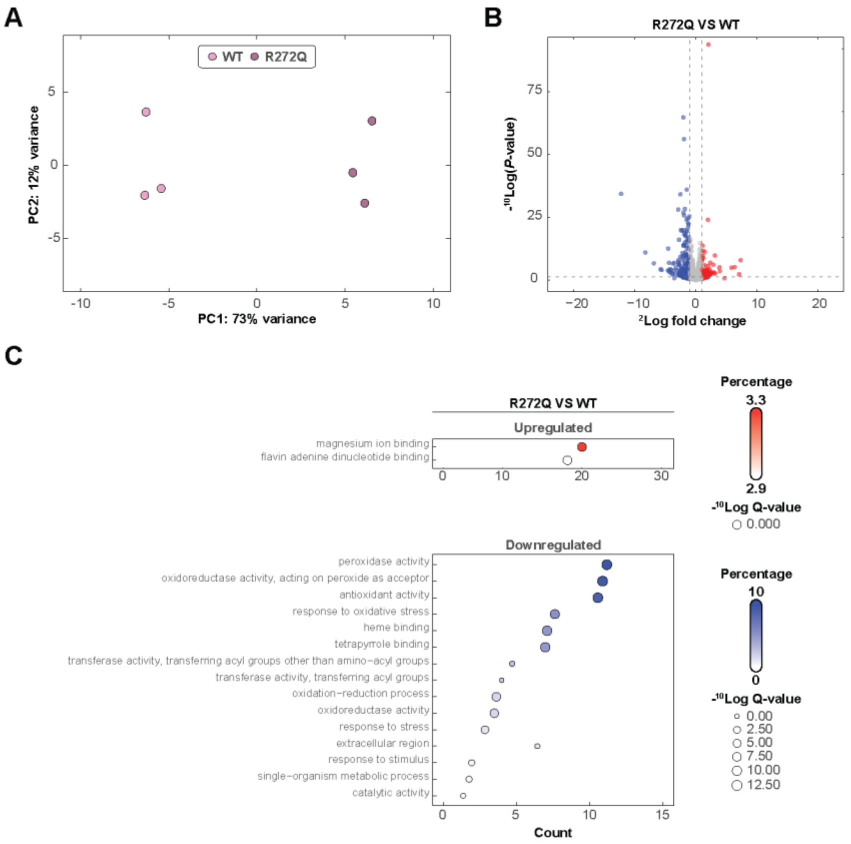


**Fig. S5. Transcriptome analysis in the MpARF1$^{R272Q}$ mutant.**

(**A**) Principal component analysis of eight-day old plants of the Mp*arf1-4* mutant complemented with MpARF1 FL$^{WT}$ and FL$^{R272Q}$ copies. Three biological replicates for both genotypes were studied. (**B**) Volcano plot showing differentially expressed genes between Mp*arf1-4* complemented with *pro*Mp*ARF1*::Mp*ARF1-FL$^{WT}$-mNG* and *pro*Mp*ARF1*::Mp*ARF1-FL$^{R272Q}$-mNG*. Genes were deemed differentially expressed when -1<$^{2}$Log(Fold Change)>1 and $P_{adj}$<.05. Genes which were significantly downregulated are blue, genes which are significantly upregulated are red. (**C**) Differentially expressed genes of fig. S5, B were used as input for gene ontology enrichment analysis. Only two terms were significantly enriched in the upregulated gene set and are shown in red, the top 15 for the downregulated gene set is shown in blue.

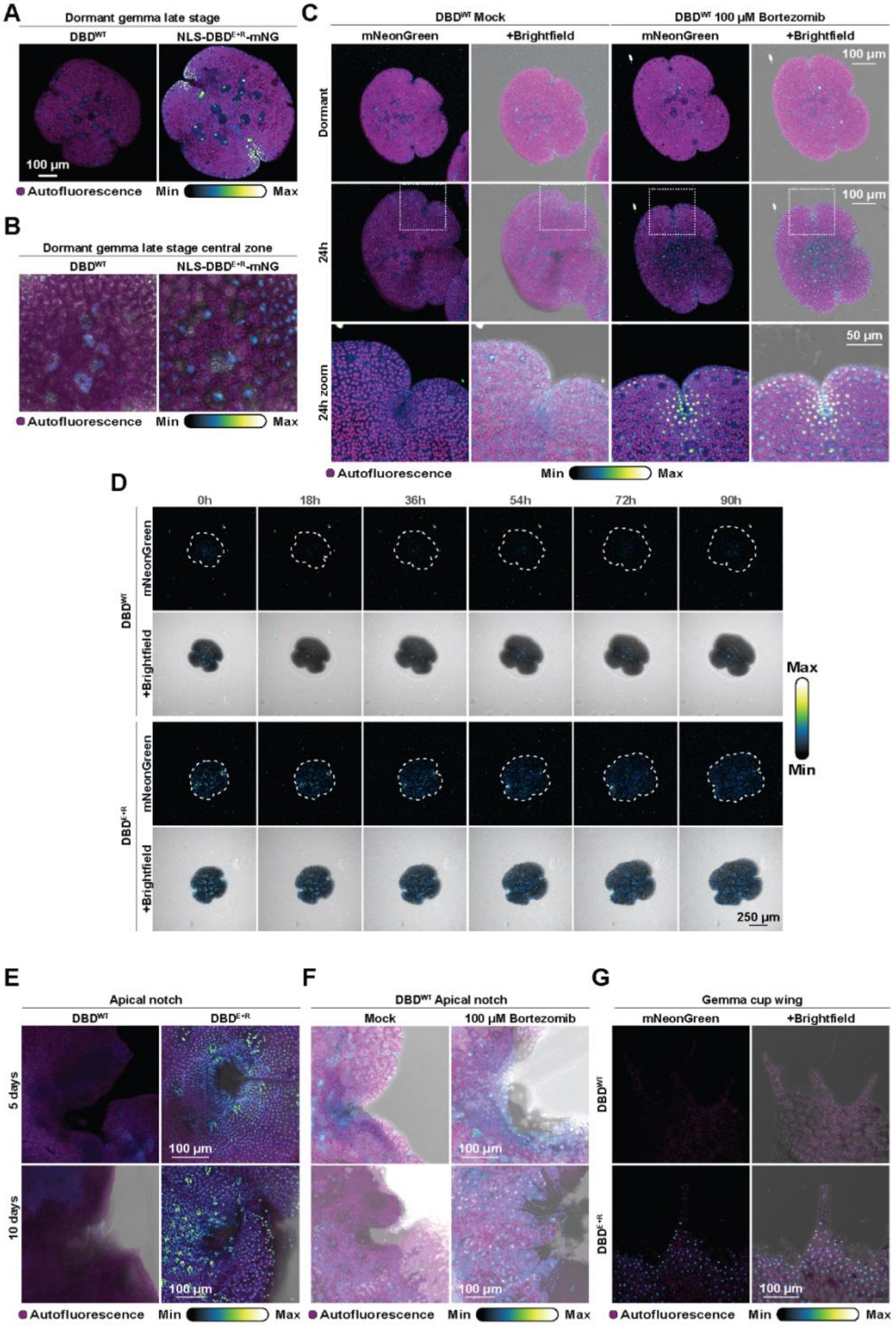


**Fig. S6. MpARF2 degradation throughout Marchantia development.**

(**A**) Micrographs depicting $DBD^{WT}$ or $DBD^{E+R}$ expressing gemmae in a late developmental stage. (**B**) Zoom in on central cells of a gemmae expressing $DBD^{WT}$ or $DBD^{E+R}$. (**C**) $DBD^{WT}$ treated with 100 µM Bortezomib (Bz) or Mock, for 24 hours. A portion of the image is outlined and enlarged below. (**D**) Timeseries experiment of gemmae expressing $DBD^{WT}$ or $DBD^{E+R}$ imaged during the early stages of gemmae germination at indicated timepoints. (**E**) Accumulation patterns of $DBD^{WT}$ or $DBD^{E+R}$ in apical notches of plants of indicated ages. (**F**) $DBD^{WT}$ accumulation patterns in apical notches treated with 100 µM Bortezomib (Bz) or Mock, for 24 hours. Plants are five days (top row) and ten days (bottom row) old, respectively. (**G**) $DBD^{WT}$ or $DBD^{E+R}$ accumulation patterns in wings of gemma cups. (A to G) Abbreviations: $DBD^{WT}$ is *pro*Mp*ARF2*::Mp*ARF2*-*DBD*$^{WT}$-*mNG* and $DBD^{E+R}$ is *pro*Mp*ARF2*::Mp*ARF2*-*DBD*$^{E+R}$-*mNG*.

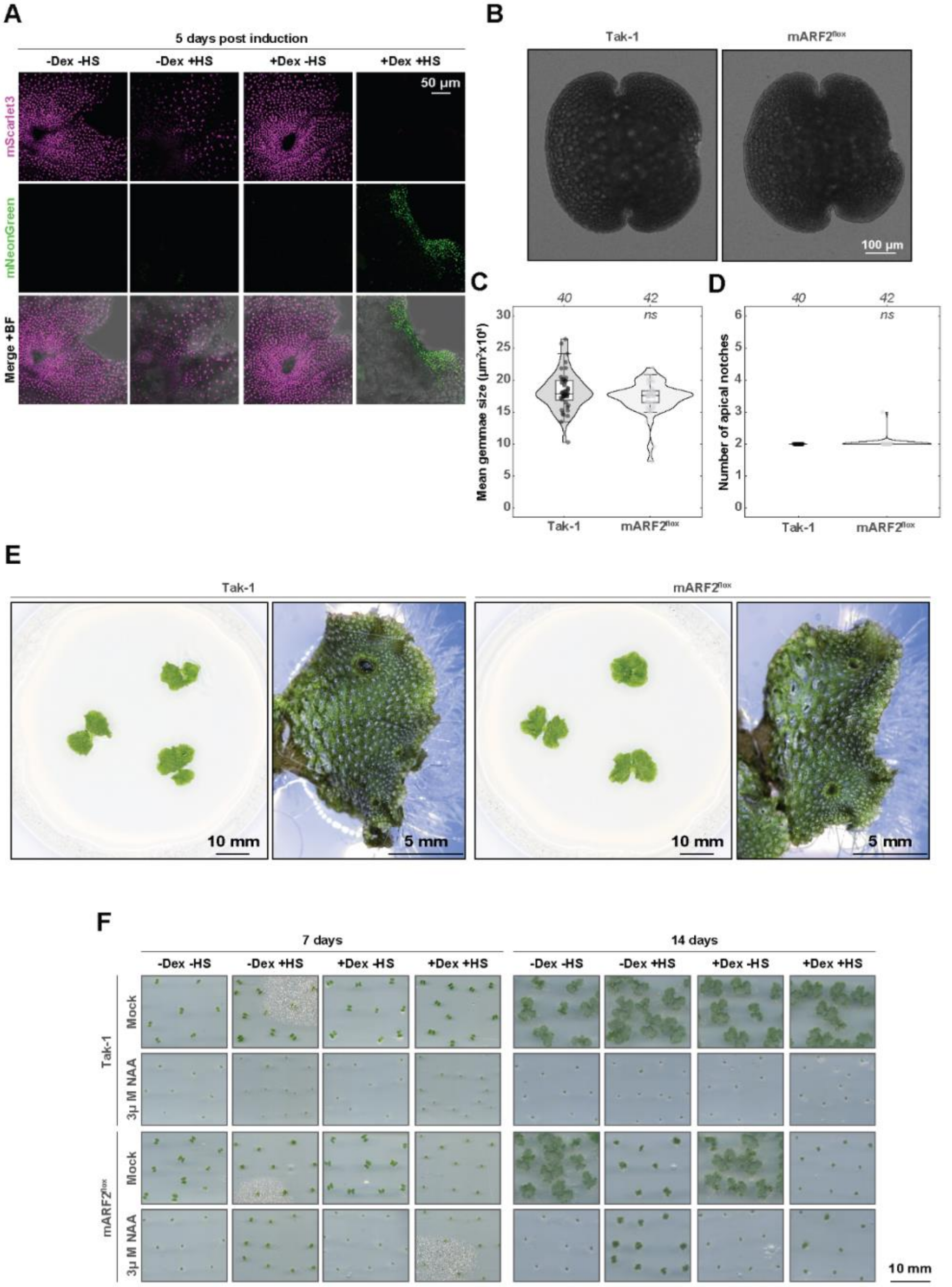


**Fig. S7. Phenotypic impact of MpARF2 accumulation.**

(**A**) Micrographs showing representative editing patterns of mARF2$^{flox}$ five days after induction with heat shock (HS) and Dexamethasone (Dex) treatment. BF; Brightfield. (**B**) Morphology of gemmae of Tak-1 and mARF2$^{flox}$ plants without induction. (**C**) Mean dormant gemmae size of Tak-1 and uninduced mARF2$^{flox}$ plants. (**D**) Average number of apical notches per gemmae of Tak-1 and uninduced mARF2$^{flox}$ plants, same experiment as (C). (**E**) Phenotypic overview of uninduced mARF2$^{flox}$ and Tak-1 plants, grown from gemmae for 14 days in normal growth conditions and on medium without treatment. (**F**) Overview of general growth of Tak-1 and mARF2$^{flox}$ plants grown on 3 µM NAA or Mock. Plants received a variety of treatments, as indicated by the different combinations of heat shock (HS) and Dexamethasone (Dex), shown above the figure. (C and D) Sample size (individual gemmae) is indicated above the figure. T-tests were used to determine statistical differences from Tak-1 ($P$<.05).

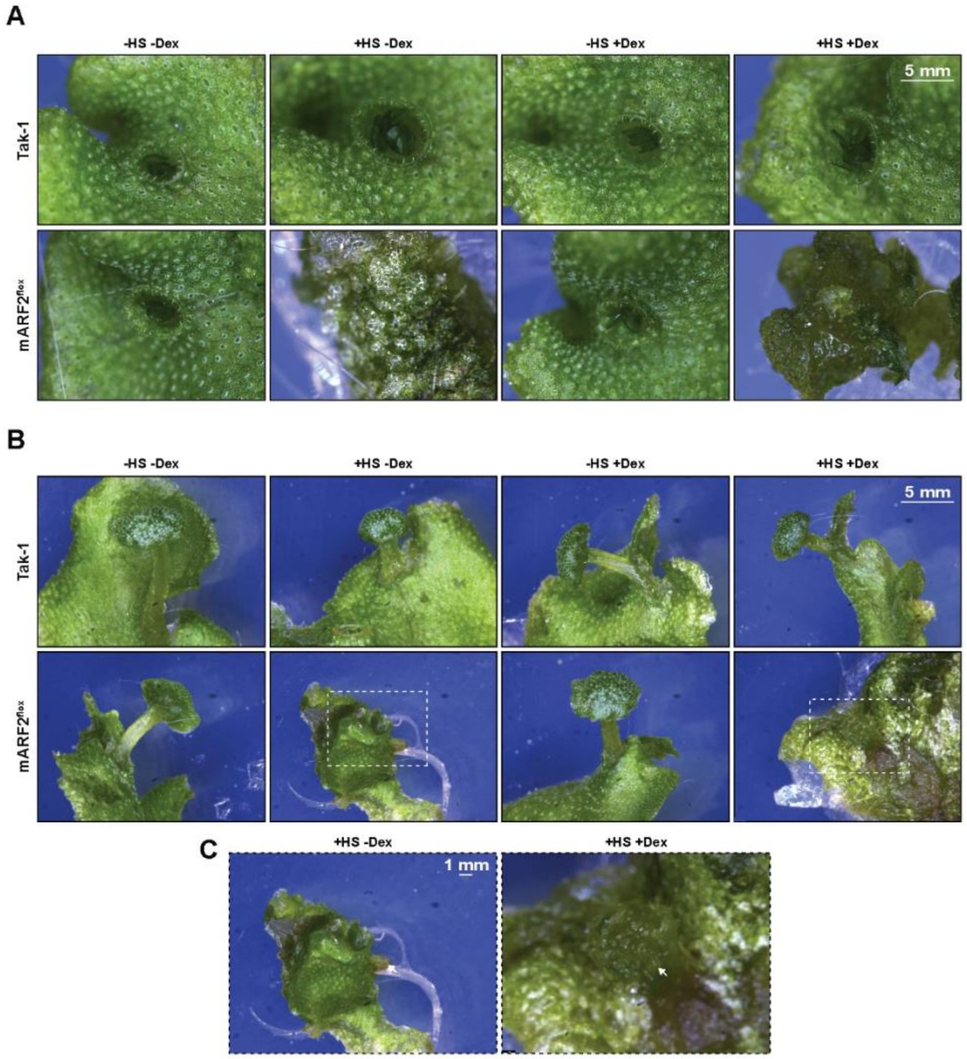


**Fig. S8. Impact of MpARF2 accumulation on gemma cup and antheridiophore development.**

(**A**) Representative images showing gemmae cup morphology of Tak-1 and mARF2$^{flox}$ plants which received the indicated HS and/or Dex treatments. (**B**) Representative images of the morphology of antheridiophores (or in mARF2$^{flox}$ +HS and +HS+Dex, putative antheridiophore-related structures) of Tak-1 and mARF2$^{flox}$ plants which received the indicated HS and/or Dex treatments. (**C**) Enlargement of the outlined regions shown in (B). (A to C) Abbreviations; heat shock (HS) and Dexamethasone (Dex).

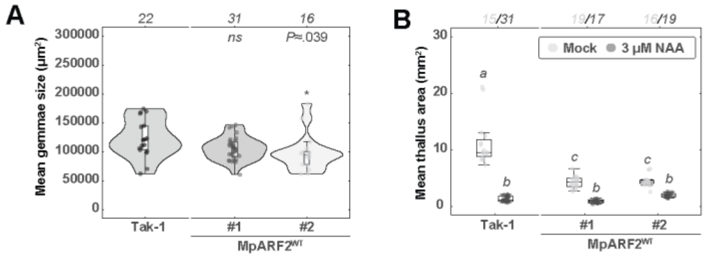


**Fig. S9. Quantification of phenotypes in MpARF2$^{WT}$ lines.**

(**A**) Mean gemma size of gemmae from Tak-1 as well as from Mp*arf2-1* mutant plants complemented with *pro*Mp*ARF2*::Mp*ARF2-FL$^{WT}$-mNG*. Size was compared per T-test ($P<.05$). (B) Projected thallus area of gemmalings grown on µM NAA or Mock for ten days. Compared are Tak-1 plants and Mp*arf2-1* mutant plants complemented with *pro*Mp*ARF2*::Mp*ARF2-FL$^{WT}$-mNG*. Mean projected thallus area was compared by Kruskal-Wallis test ($H(5)=101.38$, $P\approx2.2\times10^{-16}$) with a Dunn post hoc test ($P<.05$). Exact *P*-values can be found in the Source data.

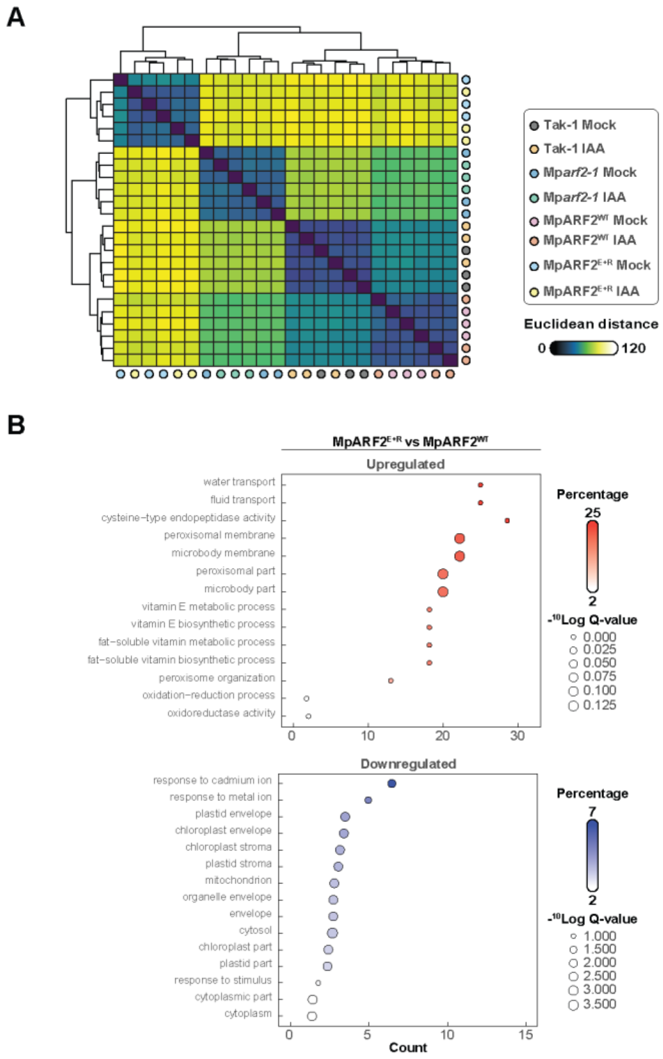


**Fig. S10. Transcriptome analysis in complemented Mp*arf2-1* mutants.**

(**A**) Hierarchical clustering of color-coded genotypes, treated with Mock or IAA, as indicated. (**B**) Gene Ontology (GO) analysis, depicting the top 15 most enriched GO terms found in the up -and downregulated gene sets of MpARF2$^{E+R}$ vs MpARF2$^{WT}$ in Mock conditions.

**Table S1.**

This table contains a description of all the plasmids used in this paper.

| no° | Backbone | Plant$^{res}$ | Bacterial$^{res}$ | Contains |
|---|---|---|---|---|
| **1** | pMpGWB100 | Hygromycin | Spectinomycin | *pro*Mp*UBE2*::*NLS*-Mp*ARF1-DBD*$^{WT}$-*mNG*-tNOS |
| **2** | pMpGWB100 | Hygromycin | Spectinomycin | *pro*Mp*UBE2*::*NLS*- Mp*ARF1-DBD*$^{R272Q}$-*mNG*-tNOS |
| **3** | pMpGWB100 | Hygromycin | Spectinomycin | *pro*Mp*UBE2*::*NLS*- Mp*ARF1-FL*$^{WT}$-*mNG*-tNOS |
| **4** | pMpGWB100 | Hygromycin | Spectinomycin | *pro*Mp*UBE2*::*NLS*- Mp*ARF1-FL*$^{R272Q}$-*mNG*-tNOS |
| **5** | pMpGWB100 | Hygromycin | Spectinomycin | *pro*Mp*UBE2*::*NLS*- Mp*ARF1-MR-mNG*-tNOS |
| **6** | pMpGWB100 | Hygromycin | Spectinomycin | *pro*Mp*UBE2*::*NLS*- Mp*ARF1-PB1-mNG*-tNOS |
| **7** | pMpGWB100 | Hygromycin | Spectinomycin | *pro*Mp*UBE2*::*NLS*- Mp*ARF1-FL*$^{WT}$-*mNG*-tNOS |
| **8** | pMpGWB100 | Hygromycin | Spectinomycin | *pro*Mp*ARF1*::*NLS*- Mp*ARF1-FL*$^{R272Q}$-*mNG*-tNOS |
| **9** | pMpGWB100 | Hygromycin | Spectinomycin | *pro*Mp*ARF2*::*NLS*- Mp*ARF1-DBD*$^{WT}$-*mNG*-tNOS |
| **10** | pMpGWB100 | Hygromycin | Spectinomycin | *pro*Mp*ARF2*::*NLS*- Mp*ARF1-DBD*$^{R272Q}$-*mNG*-tNOS |
| **11** | pMpGWB300 | Chlorsulfuron | Spectinomycin | *pro*Mp*ARF1*:: Mp*ARF1-FL*$^{WT}$-*mNG*-tNOS |
| **12** | pMpGWB300 | Chlorsulfuron | Spectinomycin | *pro*Mp*ARF1*:: Mp*ARF1-FL*$^{R272Q}$-*mNG*-tNOS |
| **13** | pMpGWB100 | Hygromycin | Spectinomycin | *pro*Mp*ARF2*::*NLS*- Mp*ARF2-DBD*$^{WT}$-*mNG*-tNOS |
| **14** | pMpGWB100 | Hygromycin | Spectinomycin | *pro*Mp*ARF2*::*NLS*- Mp*ARF2-DBD*$^{E297K+R300Q}$-*mNG*-tNOS |
| **15** | pMpGWB100 | Hygromycin | Spectinomycin | *pro*Mp*ARF2*::*NLS*- Mp*ARF2-FL*$^{WT}$-*mNG*-tNOS |
| **16** | pMpGWB100 | Hygromycin | Spectinomycin | *pro*Mp*ARF2*::*NLS*- Mp*ARF2-FL*$^{E297K+R300Q}$-*mNG*-tNOS |
| **17** | pMpGWB100 | Hygromycin | Spectinomycin | mARF2$^{flox}$ |
| **18** | pMpGWB100 | Hygromycin | Spectinomycin | *pro*Mp*ARF2*::*NLS*- Mp*ARF2-DBD*$^{A294G}$-*mNG*-tNOS |
| **19** | pMpGWB100 | Hygromycin | Spectinomycin | *pro*Mp*ARF2*::*NLS*- Mp*ARF2-DBD*$^{A295G}$-*mNG*-tNOS |
| **20** | pMpGWB100 | Hygromycin | Spectinomycin | *pro*Mp*ARF2*::*NLS*- Mp*ARF2-DBD*$^{T296N}$-*mNG*-tNOS |
| **21** | pMpGWB100 | Hygromycin | Spectinomycin | *pro*Mp*ARF2*::*NLS*- Mp*ARF2-DBD*$^{E297A}$-*mNG*-tNOS |
| **22** | pMpGWB100 | Hygromycin | Spectinomycin | *pro*Mp*ARF2*::*NLS*- Mp*ARF2-DBD*$^{K298A}$-*mNG*-tNOS |
| **23** | pMpGWB100 | Hygromycin | Spectinomycin | *pro*Mp*ARF2*::*NLS*- Mp*ARF2-DBD*$^{K298E}$-*mNG*-tNOS |
| **24** | pMpGWB100 | Hygromycin | Spectinomycin | *pro*Mp*ARF2*::*NLS*- Mp*ARF2-DBD*$^{S299A}$-*mNG*-tNOS |
| **25** | pMpGWB100 | Hygromycin | Spectinomycin | *pro*Mp*ARF2*::*NLS*- Mp*ARF2-DBD*$^{R300A}$-*mNG*-tNOS |
| **26** | pMpGWB100 | Hygromycin | Spectinomycin | *pro*Mp*ARF2*::*NLS*- Mp*ARF2-DBD*$^{F301A}$-*mNG*-tNOS |
| **27** | pMpGWB100 | Hygromycin | Spectinomycin | *pro*Mp*ARF2*::*NLS*- Mp*ARF2-DBD*$^{MpARF1\ swap}$-*mNG*-tNOS |
| **28** | pMpGWB100 | Hygromycin | Spectinomycin | *pro*Mp*ARF2*::*NLS*- Mp*ARF2-DBD*$^{MpARF1\ swap+R272Q}$-*mNG*-tNOS |

**Table S2.**

This table contains a description of the oligonucleotides used in this paper.

| Primer | Sequence (5' > 3') | Description |
|---|---|---|
| MdR329 | CCGCTGCCCATGGGGCAACG GAGAAAT | Fw introduce A294G in MpARF2 |
| MdR330 | GATTTCTCCGTTGCCCCATG GGCAGCGG | Rv introduce A294G in MpARF2 |
| MdR331 | GCTGCCCATGCGGGAACGGA GAAATCTC | Fw introduce A295G in MpARF2 |
| MdR332 | GAGATTTCTCCGTTCCCGCAT GGGCAGC | Rv introduce A295G in MpARF2 |
| MdR333 | GCCCATGCGGCAAATGAGAA ATCTCGATTC | Fw introduce T296N in MpARF2 |
| MdR334 | GAATCGAGATTTCTCATTTGC CGCATGGGC | Rv introduce T296N in MpARF2 |
| MdR335 | CTGCCCATGCGGCAGCGGAG AAATCTCGA | Fw introduce T296A in MpARF2 |
| MdR336 | TCGAGATTTCTCCGCTGCCG CATGGGCA | Rv introduce T296A in MpARF2 |
| MdR337 | CCCATGCGGCAACGGCGAAA TCTCGAT | Fw introduce E297A in MpARF2 |
| MdR338 | ATCGAGATTTCGCCGTTGCC GCATGGG | Rv introduce E297A in MpARF2 |
| MdR339 | CATGCGGCAACGGAGGCATC TCGATTC | Fw introduce K298A in MpARF2 |
| MdR340 | GAATCGAGATGCCTCCGTTG CCGCATG | Rv introduce K298A in MpARF2 |
| MdR341 | CCATGCGGCAACGGAGGAAT CTCGATT | Fw introduce K298E in MpARF2 |
| MdR342 | AATCGAGATTCCTCCGTTGCC GCATGG | Rv introduce K298E in MpARF2 |
| MdR343 | GCGGCAACGGAGAAAGCTCG ATTCTCTCTAATT | Fw introduce S299A in MpARF2 |
| MdR344 | AATTAGAGAGAATCGAGCTTT CTCCGTTGCCGC | Rv introduce S299A in MpARF2 |
| MdR345 | GCGGCAACGGAGAAATCTGC ATTCTCTCTAATTTAC | Fw introduce R300A in MpARF2 |
| MdR346 | GTAAATTAGAGAGAATGCAGA TTTCTCCGTTGCCGC | Rv introduce R300A in MpARF2 |
| MdR347 | CGGCAACGGAGAAATCTCGA GCCTCTCTAATTTAC | Fw introduce F301A in MpARF2 |
| MdR348 | GTAAATTAGAGAGGCTCGAG ATTTCTCCGTTGCCG | Rv introduce F301A in MpARF2 |
| MdR375 | CGATTGCCCTCGGTTCTTTTA TGGCTCCAAAGAAGAAGAGA AAGG | Fw SV40 to clone with hifi into *proMpARF1*::XbaI-mNG at XbaI site |
| MdR376 | ATGCTGCCGCCGCCAAGCTG TAACGGCTCAATTTCCCACAG AGAC | Rv MpARF1 DBD with hifi into *proMpARF1::XbaI-mNG at XbaI site* |

| | | |
|---|---|---|
| MdR377 | CATGCTGCCGCCGCCAAGCTGCTGCATAAATTGGCTATCATTTATAC | Rv A1 MR with hifi into *proMpARF1*::XbaI-mNG at XbaI site |
| MdR378 | ATGCTGCCGCCGCCAAGCTGGGGGCACCCCCGCTGGGCAT | Rv A1 PB1 with hifi into *proMpARF1*::XbaI-mNG at XbaI site |
| MdR379 | GCCACCAATAGCCAATTCACTATCTTTTAC | Fw A1 DBD R272Q mutation for 2 fragment assembly |
| MdR380 | GTAAAAGATAGTGAATTGGCTATTGGTGGC | Rv A1 DBD R272Q mutation for 2 fragment assembly |
| MdR432 | CCGCTGCCCATGCGGCAGCGACGAATTCTCAATTCTCTCTAATTTACAACCC | Fw swap MpARF1 R272Q mutation |
| MdR433 | GGGTTGTAAATTAGAGAGAATTGAGAATTCGTCGCTGCCGCATGGGCAGCGG | Rv swap MpARF1 R272Q mutation |
| MdR439 | GTGCCAAGCTTGCATGCCTGCAGGTCGACTACGGGGCGGATAAAAATTTTGG | Fw *pro*Mp*UBE2* to clone into pMpGWB100 XbaI |
| MdR440 | CTTCTTTGGAGCCATGGGCACCAGCACCGCCAAGCG | Rv *pro*Mp*UBE2* to clone with NLS-something into pMpGWB100 XbaI |
| MdR441 | GCGGTGCTGGTGCCCATGGCTCCAAAGAAGAAGAGAAAGG | Fw NLS SV40 to clone with *pro*Mp*UBE2* into pMpGWB100 XbaI |
| MdR442 | GGAAATTCGAGCTCGGTACCCGGGGATCCTTTACTTGTACAGCTCGTCCATGC | Rv mNG to clone into pMpGWB100 XbaI |
| MdR443 | GGGGACAAGTTTGTACAAAAAAGCAGGCTTAATGTCAGAAGCATCTTCCATCAC | Fw MpARF2 CDS (no NLS) to clone into pDONR221 linearized |
| MdR444 | ACGAAGTTATCAAGGTCGGAACTTCTGTCTAAAATGCTAG | Rv pMpARF2 to amplify pMpGWB100 XbaI to linearize and clone mARF2$^{flo}$ |
| MdR445 | GTCATAGCTGTTTCCTGTGTGAAATTGTTATCCGC | Fw pGWBO100 to amplify pMpGWB100 XbaI to linearize and clone mARF2 |
| MdR446 | ACAGAAGTTCCGACCTTGATAACTTCGTATAGCAT | Fw lox mSc3 to clone mARF2$^{flox}$ |
| MdR447 | ATGCTTCTGACATTGGATAACTTCGTATAATGTATGCTATACGAAGTTATAGC | Rv lox mSc3 to clone mARF2$^{flox}$ |
| MdR448 | ATACGAAGTTATCCAATGTCAGAAGCATCTTCCATCACTC | Fw MpARF2 CDS (no NLS) to clone mARF2$^{flox}$ |
| MdR449 | TAATGGCTGCGCGCCTGATCTAGTAACATAGATGACACCGCG | Rv MpARF2 CDS (binds tNOS, no NLS) to clone mARF2$^{flox}$ |
| MdR450 | TCTATGTTACTAGATCAGGCGCGCAGCCATTATAGC | Fw *proHSP*, to clone CRE-GR-tNOS into mARF2$^{flox}$ |
| MdR451 | GCTATGACGATCTAGTAACATAGATGACACCGCGC | Rv tNOS to clone CRE-GR-tNOS into mARF2$^{flox}$ |
| MdR452 | AAAGCAGGCTTAATGGCTCCAAAG | Fw attb+NLS to genotype plants |
| MdR453 | CATCGCCGATACCAGTAATAGTTCCC | Rv MpARF1 DBD near AD domain to genotype plants |

| | | |
|---|---|---|
| MdR454 | ATGGCTCCAAAGAAGAAGAG AAAGGTC | Fw NLS to genotype plants |
| MdR455 | GCCATCTAAGCGGGTCGACA TC | Rv MpARF1 DBD near AD domain to genotype plants |
| MdR456 | GGAATTTAGAAAGATTTCATG CG | Fw *proHSP* to seq |
| MdR457 | CTGGCATTTCTGGGGATTGC | Fw CRE to seq |
| MdR491 | GCAGCCGCCACCAATAGCCA ATTCACTATCTTTTAC | Fw R272Q MpARF1 2 fragment assembly |
| MdR492 | GCCGCCACCAATAGCCAATT CACTATCTTTTACAATCCCAG | Fw R272Q MpARF1 2 fragment assembly |
